# Dynamin-2 controls phagophore maturation

**DOI:** 10.1101/241901

**Authors:** Alejandro Martorell Riera, Cinta Iriondo Martinez, Samuel Itskanov, Janos Steffen, Brett Roach, Carla M. Koehler, Alexander M. van der Bliek

## Abstract

Autophagy involves rapid growth of phagophores through membrane addition. Newly added membranes are derived from other organelles through vesicles carrying the Atg9 protein. Membrane is delivered by fusing these vesicles with the phagophores. Atg9 is, nevertheless, not incorporated in autophagosomes. We now show that this protein is retrieved from phagophores by fission utilizing Dynamin-2 (Dnm2) as the membrane scission protein. Blocking Atg9 recycling by interfering with Dnm2 helps retain Atg9 in autophagosomes and degrades this protein by autophagy. Dnm2 colocalizes with the BAR domain protein Endophilin-B1 (EndoB1/Bif-1) when autophagy is induced, consistent with transient interactions during Atg9 retrieval. EndoB1 and Dnm2 also control the downstream fusion of phagophores to late endosomes, thus ensuring the completion of phagophores before proceeding to the next stage in the autophagy process. These data provide novel insights into the roles of membrane scission proteins during autophagy.

## Introduction

Autophagy is a ubiquitous process for disposal of damaged cytoplasmic components and recycling of proteins during starvation (Choi, Ryter et al., 2013). Much has been learned about this process by studying a series of protein complexes that perform the rapid and intricate dance to form autophagosomes, which then fuse with lysosomes for degradation (Hurley & Young, 2017, Levine & Klionsky, 2017). Autophagosomes are double membrane enclosures that fully seal off cargo destined for degradation in lysosomes (Bento, Renna et al., 2016). The membranes that form autophagosomes are derived from a variety of sources, including the plasma membrane, endosomes and the Golgi apparatus (Tooze, 2013). During the initial phases of autophagy these membranes are transported in the form of vesicles to a phagophore, which is the membrane compartments that then grows around the cargo to become a mature autophagosome once the cargo is sealed off from the cytosol. The phagophores themselves were recently shown to start off as recycling endosomes, often in close proximity to the endoplasmic reticulum (Puri, Vicinanza et al., 2018). Vesicles that bring new membrane to the phagophores also carry the Atg9 protein (Yamamoto, Kakuta et al., 2012). Atg9 is one of the few dedicated autophagy proteins with transmembrane segments. It was therefore surprising to find is retrieved when Atg9 vesicles deliver their membrane to phagophores. The mechanisms for Atg9 retrieval are currently unknown.

In this study, we focused on possible roles of Dynamin-2 (Dnm2) in autophagy. Dnm2 is one of three classic dynamins in mammalian cells (Antonny, Burd et al., 2016, Ferguson & De Camilli, 2012, Schmid & Frolov, 2011). Dnm2 is ubiquitously expressed, promoting membrane scission in a variety of different endocytic processes, including the formation of clathrin- or caveolin-coated vesicles and FEME (Renard, Simunovic et al., 2015), while dynamin-1 and dynamin-3 are primarily expressed in neurons where they act at pre- and post-synaptic densities in endocytosis (Ferguson & De Camilli, 2012). Over the years, a number of alternative functions for Dnm2 were suggested. These functions include a role in generating exocytic vesicles that bud from the TGN (Jones, Howell et al., 1998, Liu, Surka et al., 2008), in generating actin comets emanating from endocytic vesicles (Lee & De Camilli, 2002, Orth, Krueger et al., 2002), in facilitating a late stage of cytokinesis (Thompson, Skop et al., 2002), in promoting a late stage of mitochondrial fission (Lee, Westrate et al., 2016), in regenerating autolysosomes during lipid droplet breakdown in hepatocytes (Schulze, Weller et al., 2013) and in forming ATG9 containing vesicles from recycling endosomes during autophagy (Soreng, Munson et al., 2018, Takahashi, Tsotakos et al., 2016). Many of these proposed functions await further verification, but not the role of Dnm2 in endocytosis, because this has been observed in a wide range of different contexts.

All three dynamins have very similar sequences. Each one has five distinct protein domains: an N-terminal GTPase domain, a middle domain, a pleckstrin homology (PH) domain, a GTPase effector domain (GED) and a proline rich domain (PRD) (Ferguson & De Camilli, 2012). The middle domain and GED fold back on each other to form a stalk, while the PH domain interacts with membranes and the PRD binds to SH3 domains in a range of different adaptor proteins (Chappie, Acharya et al., 2010, Faelber, Posor et al., 2011, Ford, Jenni et al., 2011, Gao, von der Malsburg et al., 2010a). A large pool of dynamin proteins exists as homomeric tetramers in the cytosol, but a small fraction gathers in spots on the plasma membrane to promote membrane scission (Schmid & Frolov, 2011). Multiple contacts between adjacent dynamin molecules in those spots allow for assembly into chains, which then form multimeric spirals that wrap around the necks of budding endocytic vesicles and constrict to sever the membrane while hydrolyzing GTP (Schmid & Frolov, 2011).

One of the deciding factors for targeting dynamins to different membranes is the presence or absence of specific binding partners. Dynamins interact with PI(4,5)P_2_ at the plasma membrane (Achiriloaie, Barylko et al., 1999) and with SH3 domain containing proteins, such as amphiphysin and endophilin A during endocytosis (Simpson, Hussain et al., 1999). Besides SH3 domains, amphiphysin and endophilin A both also have BAR domains, which bind to membranes and induce curvature through their concave shapes (Daumke, Roux et al., 2014, McMahon & Boucrot, 2015). Cooperative binding interactions between dynamin and amphiphysin or endophilin A have been shown to help tubulate membranes *in vitro* and promote membrane severing *in vivo* (Farsad, Ringstad et al., 2001, Meinecke, Boucrot et al., 2013, Renard et al., 2015). Amphiphysin and endophilin A both have closely-related isoforms, amphiphysin-2 (Bin-1) and endophilin B1 (EndoB1/Bif1), that may act on other membranes (Farsad et al., 2001, Lee, Marcucci et al., 2002).

EndoB1 was proposed to interact with UVRAG as part of an autophagy protein complex built around Vps34 and Beclin-1 (Takahashi, Coppola et al., 2007), but it was also proposed to affect Atg9 vesicle formation at recycling endosomes (Takahashi et al., 2016). While studying the roles of EndoB1 during autophagy, we similarly observed colocalization of EndoB1 and Dnm2. This prompted us to further investigate a possible role of Dnm2 during autophagy. Here, we show that Dnm2 and EndoB1 promote the retrieval of Atg9 from phagophores and they control the timing of fusion to late endosomes, thus ensuring the completion of phagophores before proceeding to the next stage in the autophagy process.

## Results

### Dnm2 and EndoB1 colocalize during autophagy, but not during apoptosis

To test for Dnm2 and EndoB1 colocalization, we conducted immunofluorescence experiments with wildtype MEF cells treated with or without rapamycin to induce autophagy. These cells were permeabilized with digitonin to improve the signal from membrane bound proteins by washing out soluble cytosolic proteins (Liu, Marks et al., 1998). We observed a marked increase in Dnm2 and EndoB1 colocalization after rapamycin treatment (Fig 1A), as also determined with Mander’s correlation coefficient (Fig 1B) and with scatter plots (Fig EV1A and B). Similar results were obtained with Pearson’s correlation coefficient (not shown). To further test these interactions, we also used live cell imaging of transfected proteins with fluorescent tags as shown in Fig 1C and EV1C (quantified in Fig 1D and in Fig EV1C and D). In each of these cases, rapamycin and CCCP dramatically increased the degree of colocalization between Dnm2 and EndoB1.

**Figure 1.**
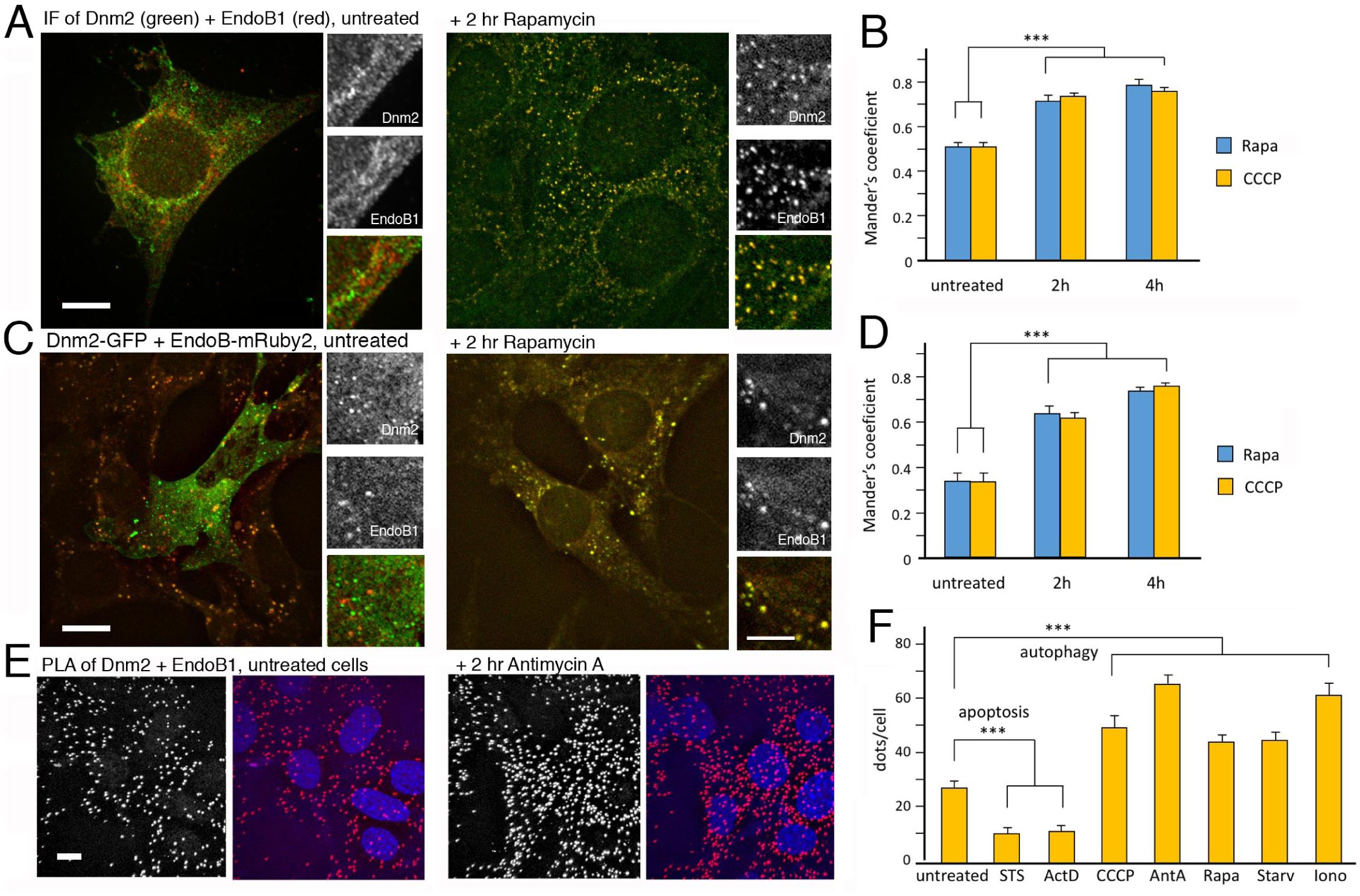
Colocalization of Dnm2 and EndoB1. A Immunofluorescence of endogenous Dnm2 (green) and EndoB1 (red) is shown without or with 2 hr rapamycin. B Mander’s coefficients for colocalization shown for untreated cells and after 2 or 4 hr with rapamycin or CCCP (50 cells were counted per experiment, SE for 3 independent experiments, unpaired Student’s t-test). C, D Similar experiments with transfected cells expressing Dnm2-GFP and EndoB1-mRuby2. E Colocalization of Dnm2 and EndoB1 tested with Proximity Ligation Assay (PLA) shown with or without 2 hr Antimycin A treatment. F Average numbers of PLA dots/cell shown for Dnm2 and EndoB1 colocalization in untreated cells, cells treated with staurosporine (STS) or actinomycin D to induce apoptosis, or with CCCP, Antimycin A, rapamycin, starvation and ionomycin to induce autophagy. Numbers of dots were counted in 50 cells/experiment (SEM, n=3, unpaired student’s t-test). Scale bar is 10 μm for whole cells and 5 μm for enlarged portions.

We further investigated colocalization of endogenous Dnm2 and EndoB1 proteins, using proximity ligation assay (PLA) because this method has a high signal to noise ratio, only giving a signal when the two proteins are within 40nm of each other. We first verified the specificity of PLA using MEF cells with homozygous deletions in chromosomal Dnm2 and EndoB1 genes, as well as MEF cells with deletions in both genes. Lack of Dnm2 and EndoB1 expression in these cell lines was confirmed with Western blots (Fig EV1E). PLA for Dnm2 and EndoB1 in these homozygous knockout cells did not yield spots (Fig EV1F and G) from which we conclude that PLA with Dnm2 and EndoB1 antibodies is highly specific. We also did not observe PLA spots when we used antibodies for Fis1, a mitochondrial outer membrane facing the cytosol, and Hsp60, a mitochondrial matrix protein, suggesting that the distance between cytosol and mitochondrial matrix is enough to preclude PLA spots (Fig EV1G). In contrast, PLA with antibodies for the bona fide partners MFF and Drp1 (Otera, Wang et al., 2010) yielded numerous spots, confirming the efficacy of this technique (Fig EV1G).

PLA was used to assess colocalization of Dnm2 and EndoB1 under a variety of autophagy and apoptosis inducing conditions. We observed dramatic increases in PLA spots with each of the autophagy inducing conditions (Fig 1E and F) in agreement with the colocalization observed by immunofluorescence and with fluorescent tagged proteins. We also expected an increase in numbers of colocalization spots during apoptosis, because Dnm2 and EndoB1 have both been connected apoptosis, albeit in different ways. Cytosolic Dnm2 promotes apoptosis in a p53 dependent manner (Fish, Schmid et al., 2000), while EndoB1 binds to Bax during apoptosis (Cuddeback, Yamaguchi et al., 2001). To our surprise, however, there was a decrease in PLA spots after treatment with apoptosis inducing chemicals. These results suggest that Dnm2 and EndoB1 have distinct functions during apoptosis and autophagy. The increase in PLA spots observed with autophagy-inducing conditions was, nevertheless, consistent with the increased colocalization observed with immunofluorescence of endogenous proteins and colocalization of transfected proteins. We therefore conclude that Dnm2 and EndoB1 colocalization increases during autophagy and decreases during apoptosis.

### Effects of Dnm2 and EndoB1 on apoptosis and mitochondrial fission

To corroborate the unexpected decrease in Dnm2 and EndoB1 interactions during apoptosis, we further tested the effects of our deletion mutants on apoptosis with two apoptosis inducing chemicals (staurosporine and actinomycin D) and three different assays (Bax recruitment to mitochondria, cytochrome c release from mitochondria and the formation of pyknotic nuclei) (Fig 2A-C). For each of these conditions, Dnm2 -/- cells showed a delay in the onset of apoptosis, while EndoB1 -/- cells showed more rapid onset of apoptosis. Dnm2-EndoB1 double knockout cells had an intermediate phenotype, which was not significantly different from wildtype. We conclude that Dnm2 has proapoptotic effects, but is not absolutely required for apoptosis, while EndoB1 is anti-apoptotic. To preclude the possibility that effects in the Dnm2 -/- cells are compensated by altered expression of the other two dynamin isoforms, we also tested apoptosis in a previously published conditional cell line in which all three dynamin genes (Dnm1, −2 and −3) can be knocked out by growing the cells with tamoxifen (Park, Shen et al., 2013). We verified this for Dnm1 and Dnm2 with Western blots (Fig EV1H). The triple knockout cells had less apoptosis than wildtype cells (Fig 2D), similar to the results obtained with Dnm2 -/- cells. We conclude that Dnm2 is pro-apoptotic, as was also shown by others (Soulet, Schmid, and Damke 2006). The anti-apoptotic effects of EndoB1 were, however, unexpected. Mutations in Dnm2 and EndoB1 also did not fully mask each other’s effects in the double knockout cells and there is less colocalization with Staurosporine and Actinomycin D (Fig 1F), suggesting that these proteins act in different pathways during apoptosis.

**Figure 2.**
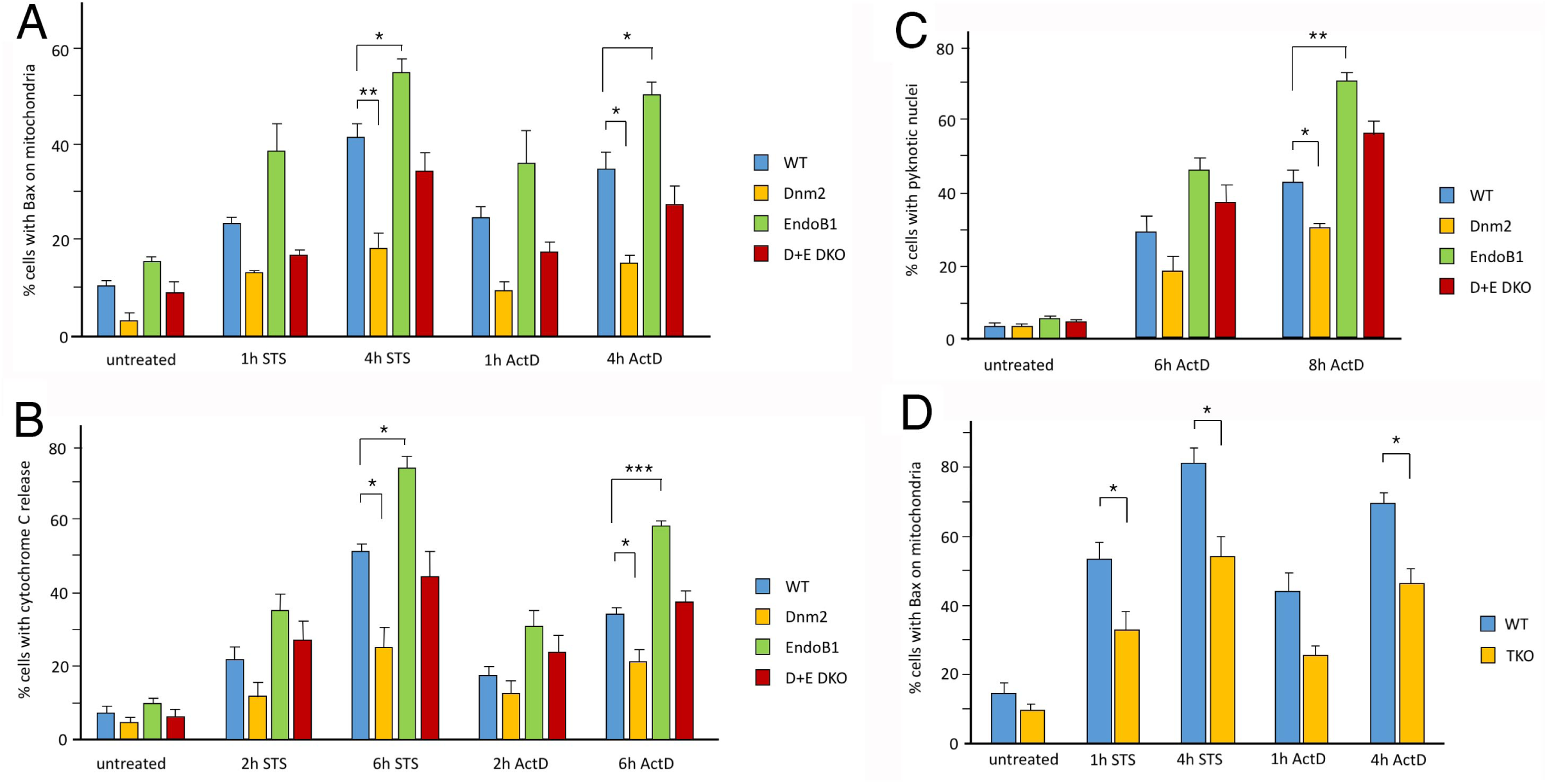
Effects of Dnm2 and EndoB1 mutations on apoptosis. A Bax translocation to mitochondria is delayed when apoptosis is induced in Dnm2 -/- cells, but not in EndoB1 -/- cells. B, C Similar effects were observed with cytochrome c release from mitochondria and with the appearance of pyknotic nuclei, which are two other indicators of apoptosis. D Bax translocation to mitochondria is delayed in cells lacking all three dynamins. Triple knockout cells (TKO) were generated by treating conditional cells (labeled as WT) with tamoxifen to induce Cre recombinase. 50 cells per experiment, SEM, n=3, unpaired Student’s t-test.

Treatment with CCCP does increase colocalization of Dnm2 and EndoB1 (Fig 1F), which would be consistent with a role in mitochondrial fission as previously suggested (Karbowski, Jeong et al., 2004, Lee et al., 2016). The potential role of Dnm2 in mitochondrial fission has, however, been contested by more recent studies, which did not show effects Dnm2 deletions on fission (Kamerkar, Kraus et al., 2018) (Fonseca, Sanchez-Guerrero et al., 2019). We tested the possible function of Dnm2 in fission by examining mitochondrial morphologies using immunofluorescence with antibodies for a mitochondrial outer membrane protein (Tom20) and a matrix protein (Hsp60). The morphologies in untreated wildtype cells, Dnm2 -/-, EndoB1 -/- and double knockout cells were not noticeably different (Fig 3A). We then treated cells with two well-established mitochondrial fission inducing chemicals (CCCP and valinomycin) (Ishihara, Jofuku et al., 2003, Yamano, Fogel et al., 2014). Both chemicals induced mitochondrial fragmentation to the same extent in mutant and wildtype cells (Fig 3A). As control, we also generated Drp1 knockout MEF cells (Fig EV2A). These cells retain highly connected mitochondria, even after treatment with CCCP, which clearly distinguishes them from Dnm2 and EndoB1 knockout cells (Fig 3B and C). Live cell imaging of mutant and wildtype cell lines also did not show differences in mitochondrial morphologies before or after treatment with CCCP (Fig EVS2B). Mitochondrial fission could still be due to increased expression of other dynamin isoforms. We tested this with the dynamin 1-3 triple knockout cells, but also did not observe differences when these cells were treated with CCCP or valinomycin (Fig 2D and E). We conclude that Dnm2 and EndoB1 are not required for CCCP or valinomycin induced fission. Colocalization of Dnm2 and EndoB1 upon treatment with CCCP (Fig 1F) could instead reflect a role in mitophagy, consistent with similar increases that were observed with other autophagy inducing treatments.

**Figure 3.**
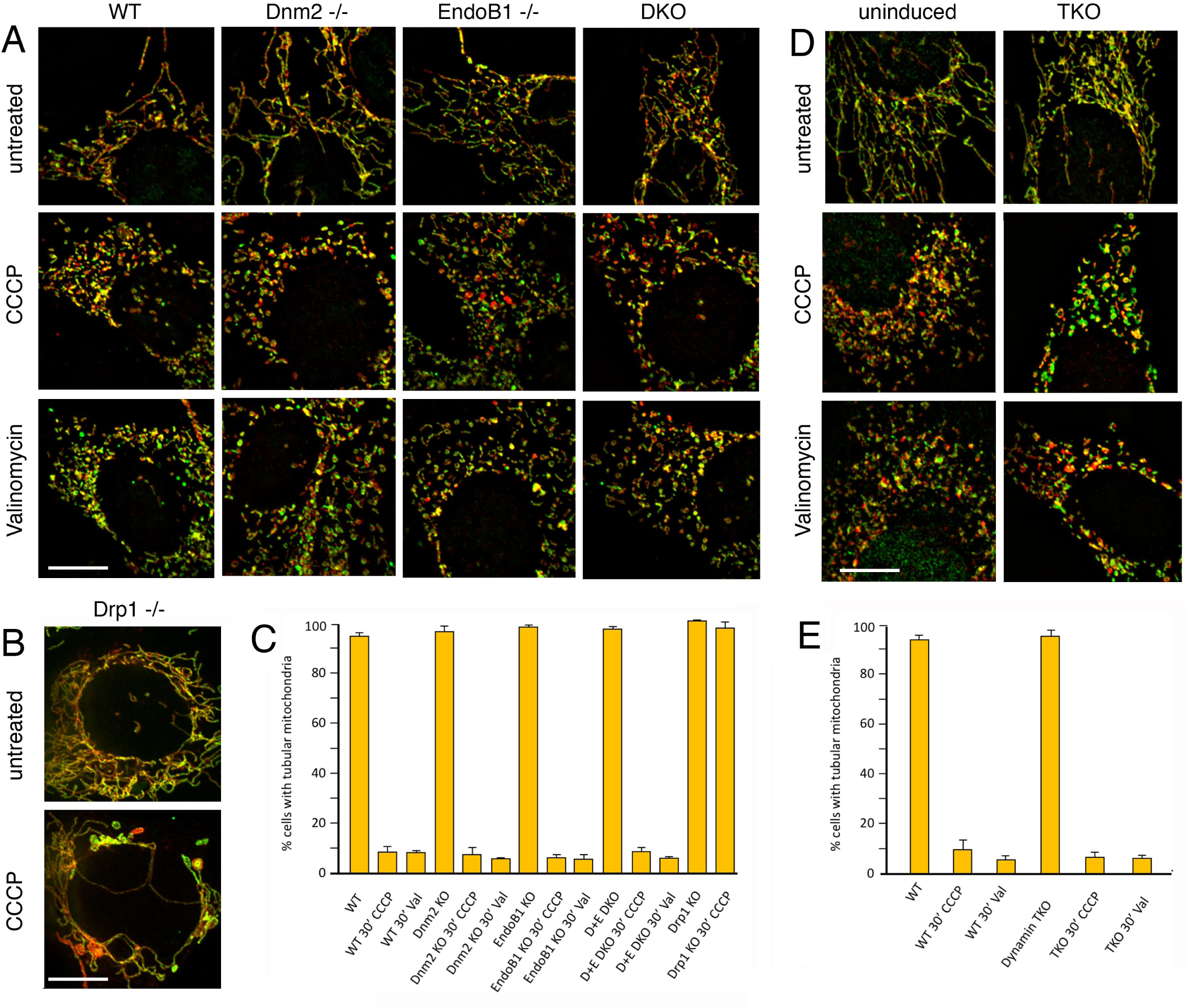
Effects of Dnm2 and EndoB1 mutations on mitochondrial fission. A Immunofluorescence of cells stained with antibodies for Tom20 (red) and Hsp60 (green) show no effects of Dnm2 or EndoB1 mutations on mitochondrial fission. The top row has images of untreated cells, the middle row has cells treated for 30 min with CCCP and the bottom row has cells treated for 30 min with valinomycin to induce mitochondrial fission. B Images of Drp1 -/- cells treated without or with CCCP. C Histogram showing that mutations in Dnm2 and EndoB1 do not prevent CCCP or valinomycin induced mitochondrial fission. D Images of cells before and after treatment with tamoxifen to induce knockout of all three dynamin genes. Cells were then treated with CCCP or valinomycin to induce mitochondrial fission. E Histogram showing that mutations in all three dynamin genes do not prevent CCCP or valinomycin induced mitochondrial fission. 50 cells/experiment, SEM, n=3, unpaired student’s t-test. Scale bar is 10 μm.

### Dnm2 colocalizes with autophagic membranes

Although the numbers of Dnm2-EndoB1 PLA spots increased when autophagy was induced, the numbers of Dnm2-EndoA1 PLA spots decreased, suggesting a shift from endocytosis to autophagy (Fig 4A). To further substantiate a possible role in autophagy, we tested colocalization with other autophagy proteins. We observed a marked increase in colocalization of PLA spots for Dnm2-EndoB1 with EGFP-LC3 when autophagy is induced (Fig 4B and C). We then tested for colocalization of PLA spots with a number of other autophagy related proteins. Results, summarized in Fig 4C, show additional increases in the fraction of Dnm2-EndoB1 PLA spots associated with two early autophagy markers (Atg2 and Fip200). We did not observe changes in colocalization with Stx17 or Rab7, which are present on mature autophagosomes, and there was a decrease in the fractions of PLA spots associated with LAMP1, which may reflect a shift away from the previously observed role in autophago-lysosomal recycling (Schulze et al., 2013). We conclude that a large fraction Dnm2-EndoB1 PLA spots are at or near sites of phagophore formation.

**Figure 4.**
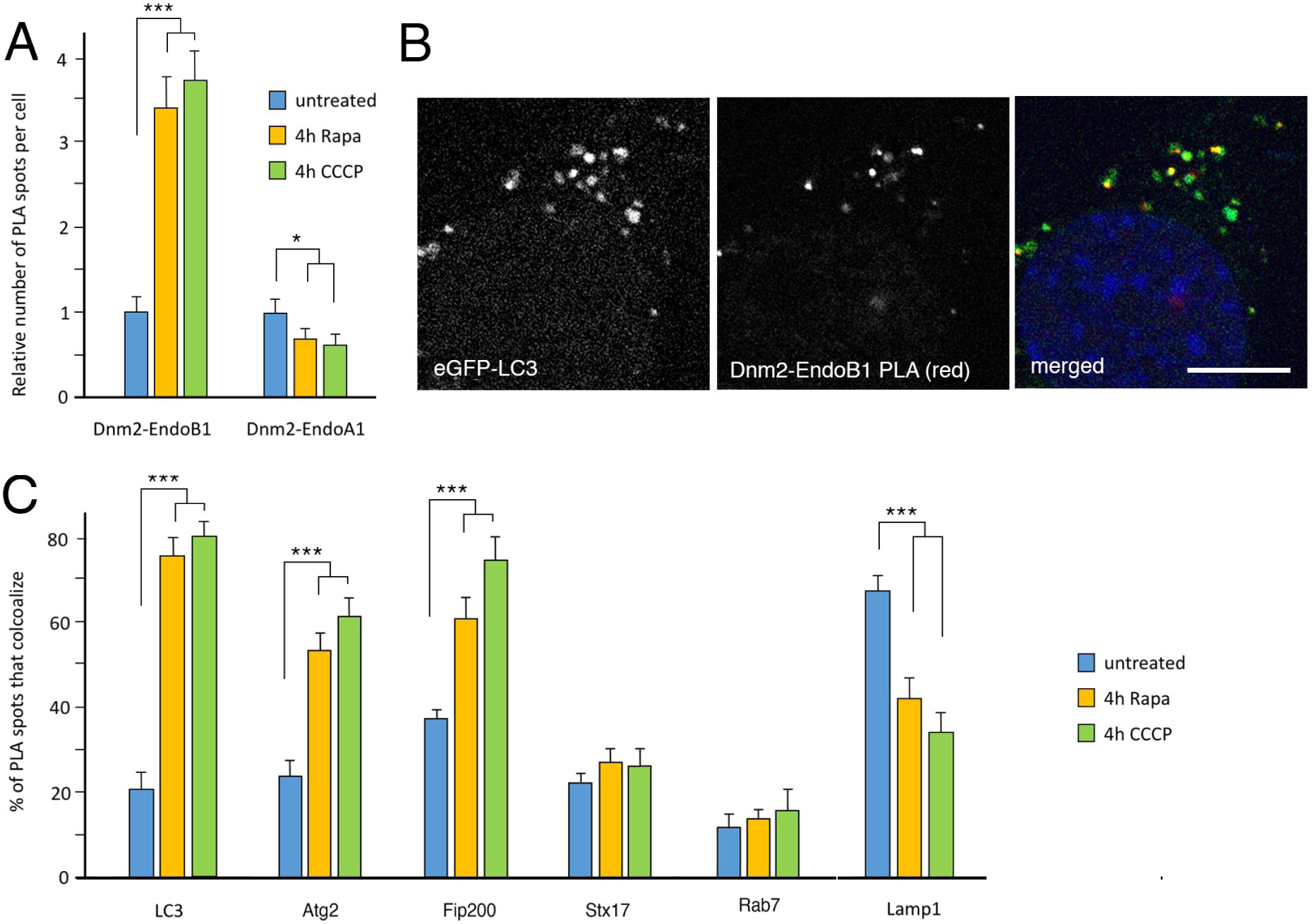
Colocalization of Dnm2 with autophagy proteins. A Relative numbers Dnm2-EndoB1 PLA spots increase, while Dnm2-EndoA1 PLA spots marginally decrease when autophagy is induced. B PLA spots for endogenous Dnm2 and EndoB1 (red) colocalize with EGFP-LC3 (green) when mitophagy is induced (4 hr CCCP). C A larger fraction of Dnm2-EndoB1 PLA spots colocalize with GFP-tagged LC3, Atg2 and Fip200 when autophagy is induced with rapamycin or CCCP, suggesting that Dnm2 is connected with autophagy. The fraction that colocalizes with GFP-tagged Lamp1 decreases and remains low for Stx17 and Rab7. For the histograms, 50 cells per experiment, SEM, n=3, unpaired Student’s t-test. Scale bar is 10 μm.

We also observed strong colocalization of Dnm2-GFP with RFP-LC3 when those tagged proteins were expressed in Dnm2 knockout cells, but not in wildtype cells (Fig EV3A). Colocalizing spots disappeared when the transfected cells were treated with the PIKfyve inhibitor YM201636 (Fig EV3A). PIKfyve is a PIP kinase associated with endosomal traffic and autophagy, converting PI(3)P into PI(3,5)P_2_, a PIP_2_ variant that can activate dynamins *in vitro* (Yarar, Surka et al., 2008) and thus could conceivably promote Dnm2 assembly and activation during autophagy. We further tested whether PIP_2_ accumulates on autophagosomes using the PLC-delta PH domain fused to GFP as a marker for PIP_2_ in Dnm2 -/- cells. This marker colocalized with LC3 spots in Dnm2 -/- cells (Fig E3B and C), suggesting that Dnm2 -/- cells have a buildup of PIP_2_ on their autophagosomes. We conclude that Dnm2 specifically localizes to phagophores when autophagy is induced and that autophagosomes accumulate Dnm2 binding sites when Dnm2 is lacking.

### Different effects of Dnm2 and EndoB1 deletions on later stages of autophagy

Since Dnm2 and EndoB1 colocalize when autophagy is induced, we tested whether both proteins contribute to this process. We first tested the effects of Dnm2 and EndoB1 deletions on CCCP induced mitophagy using fluorescence microscopy and Western blot analysis. The fluorescence images showed CCCP-induced fragmentation of mitochondria in all cells and the gradual removal of mitochondria over a period of 24 hr in WT, Dnm2-/- and DKO cells, but surprisingly not in EndoB1 single mutant cells (Fig 5A). A similar pattern was observed with Western blot analysis of the mitochondrial matrix protein Hsp60 (Fig 5B). CCCP-induced degradation of Hsp60 occurs over a period of 24 hr. This degradation is inhibited in EndoB1 -/- cells, but not in Dnm2 -/- or the DKO cells. If anything, degradation appears to be accelerated in Dnm2 -/- and DKO cells. Super resolution with structured illumination microscopy (SIM) images of EGFP-LC3 and mitochondrial DsRed shows cup-shaped phagophores surrounding mitochondria in WT, Dnm2 -/- and DKO cells (Fig 5C). These phagophores are indistinguishable from wildtype. In contrast, there were no cup shaped phagophores in EndoB1 knockout cells. EGFP-LC3 did form aggregates but those appeared to be separate from mitochondria. Together, these results show that an EndoB1 deletion inhibits mitophagy, but a Dnm2 deletion does not. Suppressive effects of the Dnm2 deletion on the EndoB1 deletion suggests that these proteins act in the same pathway but with opposite effects.

**Figure 5.**
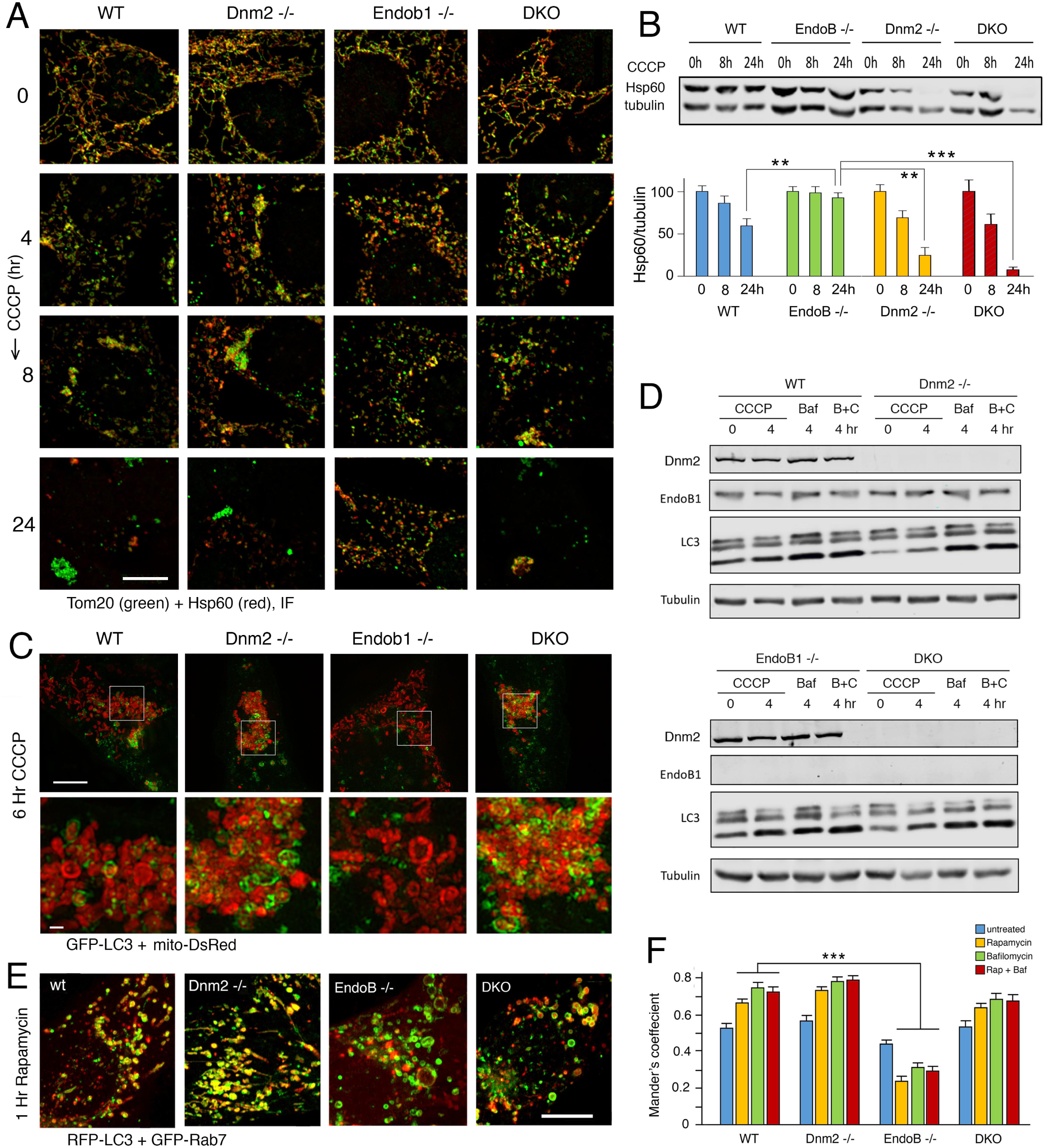
Effects of Dnm2 and EndoB1 mutations on autophagy. A Effects of CCCP on mitophagy in WT, Dnm2 -/-, EndoB1 -/- and DKO cells observed with immunofluorescence microscopy using Tom20 (red) and Hsp60 (green) antibodies. At 24 hr, most mitochondria are degraded, except in EndoB1 -/- cells, where mitophagy is inhibited. Scale bar is 10 μm. B Western blots showing the effects of CCCP on Hsp60 levels in the mutant cells. The bottom panel shows average intensities of Hsp60 bands in four independent experiments, relative to tubulin levels and normalized at 100% for 0hr time point. C Super-resolution images (SIM) show the formation of LC3-membranes (green) encapsulating mitochondria (red) in cells treated with CCCP. These structures are observed in WT, Dnm2 -/- and DKO cells, but not in EndoB1 -/- cells. Cells were treated for 6 hr with 20 μM CCCP. Mitochondria were detected with mitoDsRed and autophagic membranes were detected with EGFP-LC3. Scale bar is 10 μm for top panels and 1 μm for the enlargements. D Western blots showing the effects of CCCP and Bafilomycin A treatments on LC3 lipidation in wildtype, Dnm2 -/-, EndoB1 -/- and DKO cells. E Effects of Dnm2 and EndoB1 deletions on colocalization of RFP-LC3 and GFP-Rab7. Autophagy was induced for 1 hr with Rapamycin, Scale bar is 10 μm. F Reduced levels of LC3 and Rab7 colocalization expressed as Mander’s coefficient for untreated cells and for cells treated with Rapamycin, Bafilomycin or both. There were 50 cells per experiment, SEM, n=3, unpaired Student’s t-test.

The stage of inhibition by EndoB1 was further investigated with Western blot analysis of LC3. Lipidation of LC3 increased in EndoB1 -/- cells that were treated with CCCP, but it did not increase further with Bafilomycin, suggesting that lipidation was not affected by the EndoB1 deletion but autophagic flux was impaired. As with the mitophagy data, flux appeared to be unaffected in Dnm2 -/- and DKO cells, suggesting that the EndoB1 deletion blocks an intermediate stage of autophagy, but this block is suppressed by the Dnm2 deletion (Fig 5D, Fig EV3D). Similar effects were observed with other autophagy inducing treatments, but not with starvation. Autophagic flux induced by starvation was not impaired in EndoB1 -/- cells, because the amounts of lipidated LC3 was similarly increased in Bafilomycin treated WT, Dnm2 -/-, EndoB -/- and DKO cells (Fig EV3E).

To further investigate the stage of inhibition in EndoB -/- cells, we determined the degree of LC3 and Rab7 colocalization with fluorescence microscopy. LC3 and Rab7 colocalization was consistently reduced in EndoB -/- cells, but not in Dnm2 -/- or DKO cells (Fig 5E and F), suggesting that CCCP-induced lipidation in EndoB1 -/- cells was not enough to further progression along the autophagy pathway. These results suggest that the EndoB1 -/- deletion impairs autophagy at a stage between LC3 lipidation and fusion with Rab7 containing compartments. This impairment is reversed by a deletion in Dnm2, suggesting that the two proteins act in the same pathway. We conclude that Dnm2 can inhibit a late stage of autophagy, but this inhibition is relieved by the presence of EndoB1.

### Early stage effects of Dnm2 on Atg9 retrieval from phagophores

To determine what specific roles Dnm2 might play in autophagy, we monitored RFP-tagged Dnm2 spots at or near phagophores using time lapse images of live cells. Phagophores were labeled with BFP-LC3 and incoming vesicles that add membrane to phagophores were labeled with GFP-Atg9. We observed RFP-tagged Dnm2 spots that appeared and disappeared between BFP-LC3 labeled phagophores and GFP-Atg9 labeled organelles that approached each other and then separated within the span of 1 second. An example is shown in Fig 6A and a second example is shown in Fig EV4A. The frequency of observed events in each video was low (1 per 3 videos in untreated cells) due to limitations of the imaging system (50 images of a portion of a cell in one plane at 350 msec intervals), but an extrapolation to whole cells suggests a rate in the range of 25 events/cell/minute, which would be consistent with a role in autophagy. Because of photo-bleaching and the rapid movements of organelles, we were unable to determine whether multiple fission and fusion events occurred at the same spots on growing phagophores or if they were independent. The sequence of events within a single cycle are, nevertheless, suggestive of “kiss and run” fission and fusion, which is a well-documented dynamin-dependent process that occurs at the plasma membrane in chromaffin cells and at neuronal synapses (Alabi & Tsien, 2013, Holroyd, Lang et al., 2002). “Kiss and run” fission and fusion at the plasma membrane can be inhibited with the dynamin inhibitor Dynole 34-2 and stimulated with the dynamin activator Ryngo 1-23 (Jackson, Papadopulos et al., 2015, Lasic, Stenovec et al., 2017). We tested the effects of these chemicals on appearance and disappearance of RFP-tagged Dnm2 spots wedged between BFP-LC3 and GFP-Atg9 labeled organelles (Fig 6B). We observed a marked reduction in the numbers of these Dnm2 spots with Dynole 34-2 and an increase with Ryngo 1-23 consistent with an active role at this stage of phagophore formation.

**Figure 6.**
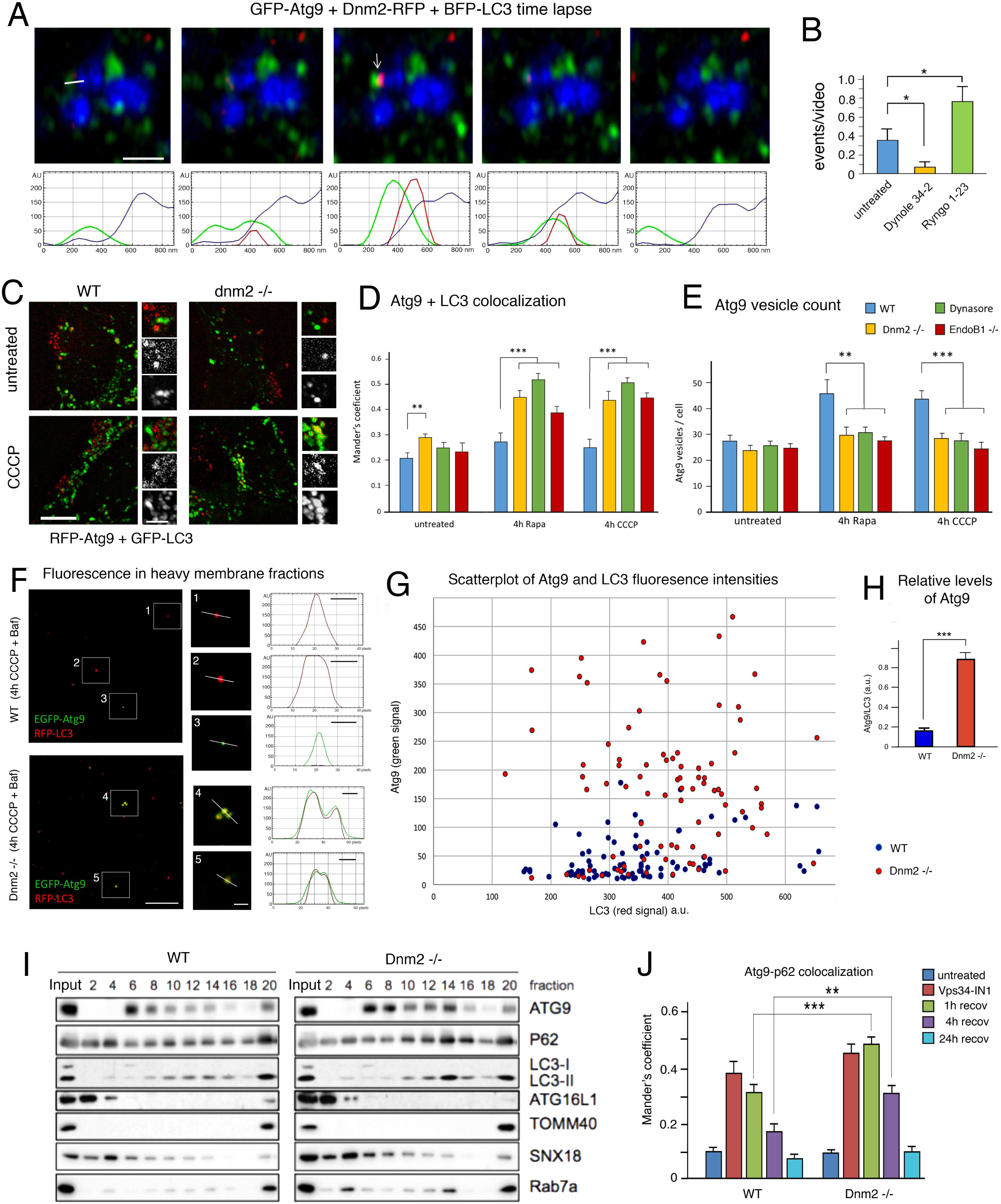
Effects of Dnm2 on Atg9 localization. A Time lapse images of GFP-Atg9, Dnm2-RFP and BFP-LC3. Images were taken at 350 msec intervals. An arrow in the middle panel points to a potential scission event. The bar in the first panel shows the line used for the intensity tracings shown in the bottom panels. Scale bar is 1 μm. B Mean frequencies of Dnm2-RFP spots transiently wedged between GFP-Atg9 and BFP-LC3 spots in untreated cells and cells treated with Dynole 34-2 or Ryngo 1-23 (n = 20, 17 and 22, SEM, unpaired T test). The frequencies were low because only a small part of a cell was observed for a limited time (50 frames), but their transient nature and the opposing effects of a dynamin inhibiter and an activator suggest a functional connection with Atg9 and LC3 fission and fusion events. C RFP-Atg9 colocalizes with EGFP-LC3 in Dnm2 -/- cells treated with CCCP, but not in wild type cells. Scale bar is 10 μm for whole cells and 3 μm for enlarged portions. D Histogram showing increased colocalization of RFP-ATG9 and GFP-LC3 in Dnm2 -/- cells, dynasore-treated WT cells and EndoB1 -/- cells after incubation with rapamycin or CCCP. E The number of Atg9 vesicles (RFP spots without GFP label) decreases when Dnm2 -/- cells, dynasore-treated WT cells and EndoB1 -/- cells are incubated with rapamycin or CCCP. For panels D and E, 50 cells were counted per experiment, n= 3, SEM, unpaired students’ t-test. F Images of autophagosome containing fractions from WT and Dnm2 -/- cells transfected with EGFP-Atg9 and RFP-LC3 followed by 4hr treatments with CCCP and Bafilomycin to induce mitophagy and to accumulate autophagosomes. Autophagosomes were enriched by differential centrifugation. Enlargements show individual spots with line tracings to plot fluorescence intensities. Scale bar is 10 μm in the overview images and 2 μm in the enlargements. The x-axis in the fluorescence intensity plots show numbers of pixels (24 pixel/μm) and the bars show 500 nm. G Scatter plot of peak fluorescence intensities for individual spots, as shown in panel F (arbitrary units). H Histogram of the peak fluorescence intensities of Atg9 in LC3 spots relative to the peak intensities of Atg9 in spots without LC3. Bar is SE, with Student’s t-test for 3 independent experiments. 30 spots for each condition were measured in each experiment. I Subcellular fractionation of homogenates from WT and Dnm -/- MEF cells, which were treated for 4 hr with Rapamycin. Membranous organelles were separated with a 1-22% Ficoll density gradient and analyzed with Western blots. The bottom fraction (#20) contains mitochondria and other dense organelles such as mitophagosomes, autophagolysosomes or lysosomes. Atg9 is normally present in light fractions (fraction #6), but partially shifts to an intermediate density (fraction #14) in Dnm2 -/- cells, along with some p62, LC3-II and Rab7a. J Delayed dissociation of Atg9 and p62 in Dnm2 -/- cells. A buildup of Atg9 and p62 was induced for 16h with the Vps34 inhibitor Vps34-IN1. This was followed by recovery through washout of the inhibitor for the indicated lengths of time. Association and dissociation of Atg9 and p62 was monitored by immunofluorescence microscopy of endogenous proteins in WT and Dnm2 -/- HeLa cells.

The Atg9 protein, which helps bring membrane to growing phagophores, is not normally incorporated in mature autophagosomes, suggesting that this protein is retrieved by a recycling mechanism. We tested possible effects of the Dnm2 deletion on the retrieval of Atg9 from autophagic membranes, because this process would involve membrane scission at phagophores. We tested whether the block in retrieval could be detected with fluorescent proteins. Dnm2 and wildtype cells were transfected with RFP-tagged Atg9 and EGFP-LC3 and fixed for microscopy. We observed well-defined colocalization of Atg9 and LC3 in Dnm2 cells, but not in wildtype cells (Fig 6C), as confirmed with Mander’s coefficients (Fig 6D). There was also a concomitant drop in the numbers of spots that were only labeled with Atg9, but not with LC3 (Fig 6E), consistent with the idea that fusion with phagophores depletes the supply of Atg9 vesicles. Similar changes in Atg9 colocalization with LC3 were observed with the dynamin inhibitor dynasore (Fig 6D and E), in live cells (Fig EV4B and C), in WT cells transfected with Dnm2 siRNA (Fig EV4D-F) and in a Dnm2 -/- HeLa cell line (Fig EV4G-I), which strengthens the conclusion that Dnm2 is needed to prevent colocalization of Atg9 and LC3 in autophagosomes. In parallel, we determined whether the EndoB1 deletion also affected Atg9 and LC3 colocalization or the numbers of separate Atg9 vesicles (Fig 6D and E). The patterns that we observed with EndoB1 -/- cells were similar to those in Dnm2 -/- cells showing that Dnm2 and EndoB1 both help prevent colocalization of Atg9 and LC3 during autophagosome maturation. This cooperative effect contrasts with the antagonism between Dnm2 and EndoB1 during the transition from phagophore to autophagosome, as described in the previous section.

To further test whether Atg9 and LC3 are present in the same compartments in Dnm2 -/- cells, we imaged individual autophagosomes in crude biochemical isolates as described (Gao, Kang et al., 2010b). MEF cells were transfected with EGFP-Atg9 and RFP-LC3 followed by 4hr treatments with CCCP and Bafilomycin to induce mitophagy and to accumulate autophagosomes. These autophagosomes were enriched by differential centrifugation and washed twice in 1.0M NaCl to dissociate clusters of vesicles that were potentially held together by ionic interactions. Samples of the resulting fractions were mounted on slides and examined by fluorescence microscopy.

Extracts from WT cells showed fluorescent spots with LC3 but little or no Atg9, presumably corresponding to mature autophagosomes (Fig 6F). An occasional spot with Atg9 but not LC3 was also observed, possibly corresponding to an Atg9 vesicle or organelle. In contrast, extracts from Dnm2 -/- cells invariably showed spots with overlapping LC3 and Atg9 fluorescence (Fig 6F). Examples of line tracings for selected spots are shown in Fig 6F, while a third example for Dnm2 -/- cells is shown in Fig EV4J-L. Tracings of LC3 and Atg9 fluorescence intensities were closely matched in Dnm2 -/- extracts, suggesting that membranes containing LC3 and Atg9 were fully merged. A scatterplot of peak intensities for different spots with LC3 and Atg9 shows that a large number of these spots carry both proteins in Dnm2 -/- extracts, but not in WT extracts (Fig 6G). A histogram of these peak intensities for three independent experiments shows that the distribution is significantly altered in Dnm2 -/- cells (Fig 6H). We conclude that Atg9 and LC3 containing membrane are merged in Dnm2 -/- cells, but not in WT cells.

Next, we used subcellular fractionation as a biochemical test for possible aberrant Atg9-containing organelles in Dnm2 -/- cells. The distribution of Atg9-containing organelles was analyzed with Ficoll density gradients as previously described (Orsi, Razi et al., 2012). Autophagy was induced in WT and Dnm2 -/- MEF cells with Rapamycin, instead of CCCP, because the mitophagosomes that are formed with CCCP could not be separated from intact mitochondria in Ficoll gradients, while Rapamycin also induces the formation of less dense autophagosome related structures that can be separated with Ficoll gradients. Atg9 is normally present in a light membrane fraction (fraction #6 in Fig 6I), but some protein is observed in the heavy membrane fractions of induced cells (#20). Other autophagy proteins, such as p62 and LC3-II, are also found in this fraction along with mitochondrial (TOMM40) and endo/lysosomal (Rab7a) proteins. Strikingly, however, Dnm2 -/- cells show an additional accumulation of certain proteins (Atg9, p62, LC3-II and Rab7a) in a novel fraction (fraction #14 in Fig 6I). This fraction does not contain Atg16L1, which acts at an early stage to promote LC3 lipidation, nor does it contain SNX18, which was previously suggested to act together with Dnm2 in Atg9 vesicle formation at early endosomes (Soreng et al., 2018). These results suggest that fraction 14 contains aberrant late stage autophagosomes with Atg9, LC3-II, Rab7a and p62. The presence of Atg9 in fraction #14 is remarkable, because this protein would not normally be present at a late stage, when Atg16L1 is removed and the more mature organelles fuse with Rab7a containing compartments.

We used a pulse-chase approach to investigate temporal aspects of autophagy affected by the Dnm2 deletion. LC3 lipidation was blocked for 16h with a Vps34 inhibitor (Vps34-IN1) followed by washout and recovery for increasing lengths of time. Atg9 and p62 colocalization were monitored by immunofluorescence in WT and Dnm2 -/- HeLa cells (these cells were used here, because Atg9 immunofluorescence was more specific with human cells than with mouse cells). We observed increased colocalization of Atg9 and p62 in WT and in Dnm2 -/- cells after 16h treatment with Vps34-IN1 (Fig 6J), consistent with clustering of Atg9 vesicles near stalled phagophores. Colocalization was gradually reduced over a 24hr period after Vps34-IN1 was washed out. This recovery was, however, significantly delayed in Dnm2 -/- cells (Fig 6J), consistent with impaired Atg9 retrieval.

We conclude that Dnm2 contributes to the retrieval of Atg9 from phagophores. Live cell imaging shows Dnm2 in transient spots between Atg9 and LC3 containing compartments. Mutations in Dnm2 shift the balance towards merging of Atg9 containing membranes with the phagophores and the formation of aberrant autophagic organelles. A role in Atg9 retrieval from phagophores conflicts with previously proposed functions in the formation of Atg9 vesicles at early endosomes (Takahashi et al., 2016), because inhibition there would slow down autophagy and lead to an accumulation of Atg9 in other compartments. The increased colocalization of Atg9 and LC3 that we observed both in Dnm2 and EndoB1 -/- cells, suggest that these proteins act together to promote Atg9 retrieval during the early stages of autophagy, while the inhibitory effects of Dnm2 that are unmasked with the EndoB1 deletion (see previous section) occur at a later stage.

### Downstream effects of the Dnm2 deletion on autophagy

To track the downstream events affected by mutations in Dnm2 during autophagy, we transfected cells with RFP-GFP-LC3, which is visible as yellow spots in the cytosol and red spots when autophagosomes fuse with lysosomes, because acidification quenches GFP. Our results show that Dnm2 -/- cells have larger numbers of red fluorescence spots than wildtype cells after treatment with CCCP (Fig 7A and B). Autophagosomes in Dnm2 -/- cells progress more rapidly towards fusion with lysosomes than autophagosomes in wildtype cells treated with CCCP (Fig 7C). Increased rates of fusion to lysosomes were also detected through the merging of RFP-LC3 with the lysosomal marker GFP-Lamp1 (Fig 7D).

**Figure 7.**
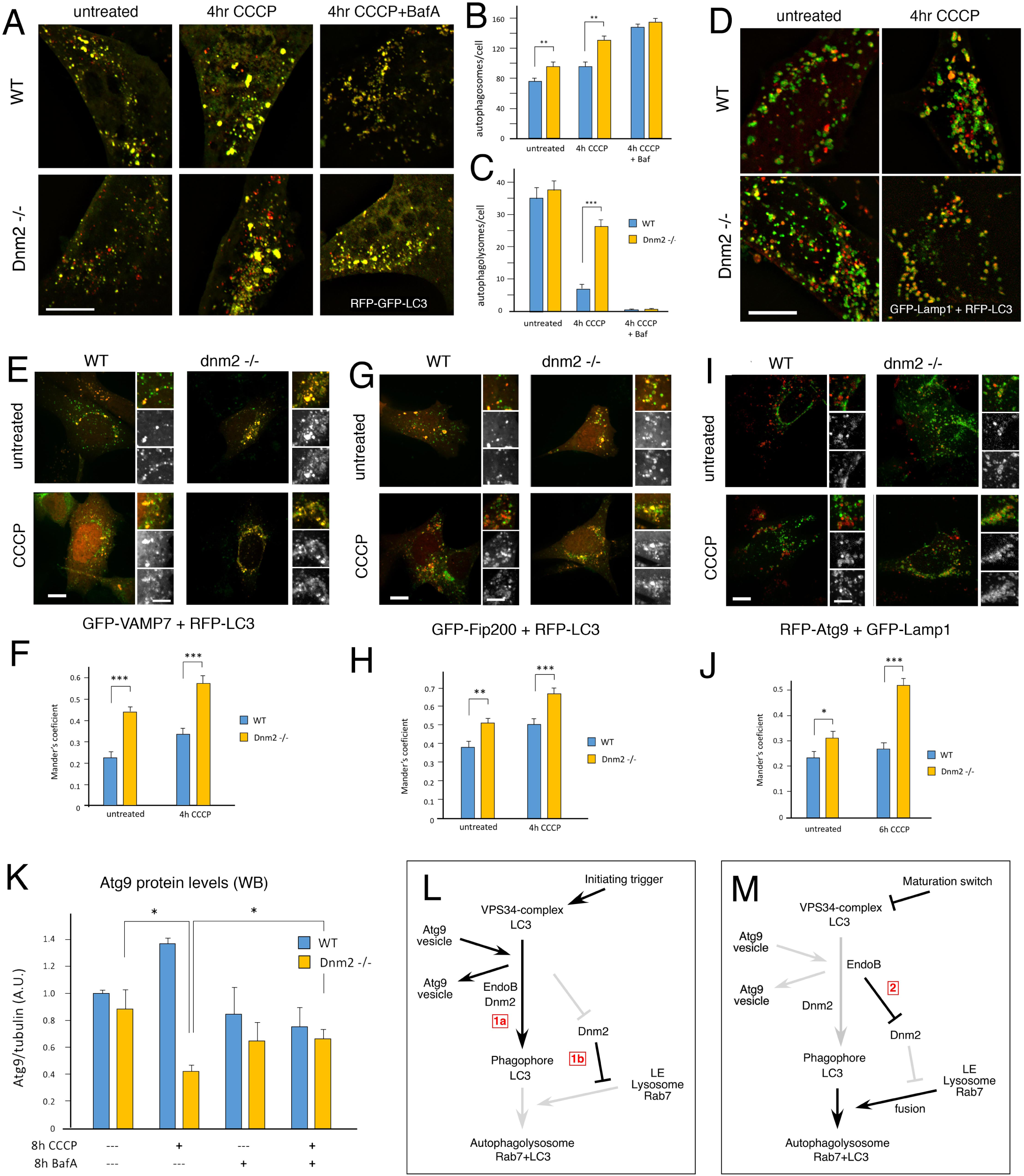
Effects of Dnm2 on Atg9 retrieval from phagophores. A Tracking autophagy with RFP-GFP-LC3 marker shows transfer of LC3 to autophagolysomes, visible as red spots due to acidic quenching of GFP. B Numbers of phagophores or autophagosomes per cells, detected as yellow spots are increased in Dnm2 -/- cells. C Numbers of autophagolysosomes, detected as red spots, are dramatically increased in Dnm2 -/- cells treated with CCCP. D Colocalization of RFP-LC3 with GFP-Lamp1 is similarly increased in Dnm2 -/- cells. E, F Colocalization of GFP-VAMP7 with RFP-LC3 is increased in Dnm2 -/- cells and increases more after CCCP treatment. G, H Colocalization of GFP-Fip200 with RFP-LC3 is increased in Dnm2 -/- cells and increases more after CCCP treatment. I, J There is also increased colocalization of RFP-Atg9 and GFP-LAMP1 after treatment with CCCP. Cells were incubated with pepstatin and E64d to prevent Atg9 digestion in lysosomes. For panels B, C, F, H and J, 50 cells were counted per experiment, n= 3, unpaired students’ t-test. Scale bar is 10 μm for whole cells and 5 μm for enlarged portions. K Quantification of Atg9 protein levels in wildtype and Dnm2 -/- cells. Levels decrease in Dnm2 -/- cells after treatment with CCCP, but this decrease is partially prevented by Bafilomycin A, suggesting that Atg9 is degraded by autophagy in Dnm2 -/- cells. Band intensities were determined with Licor software, n = 4, unpaired Students’ t-test. L Genetic representation of the proposed roles for Dnm2 and EndoB1 during autophagy. The initial stages of phagophore formation during which EndoB1, as part of the Vps34 complex, promotes the role of Dnm2 in the retrieval of Atg9 through fission from phagophores (1a). At this stage, Dnm2 also plays an inhibitory role, preventing fusion to late endosomes or lysosomes (1b), so that further maturation of the phagophore only occurs when growth of the phagophore is complete. M A later stage during which EndoB1 switches from a positive to a negative function, suppressing the inhibitory effects of Dnm2 on fusion to late endosomes and lysosomes (2).

We reasoned that the SNARE proteins carried along with Atg9 vesicles could be retrieved by a Dnm2-dependent process and that this retrieval would be disrupted in Dnm2 -/- cells. We first tested colocalization of VAMP7, which is a v-SNARE that has been localized to late endosomes but is also associated with Atg9 vesicles (Aoyagi, Itakura et al., 2018, Yu, Chen et al., 2017). Our results show increased colocalization with LC3 in Dnm2 -/- cells, and even more so after induction of mitophagy with CCCP (Fig. 7E and F). We could similarly detect increased colocalization of the Atg9 vesicle tether Fip200 with LC3 (Fig. 7G and H). Although this protein is not incorporated in phagophores and it is not membrane anchored, there could still be an increased association with LC3 due to the mistargeting of Atg9 protein to phagophores. We then tested colocalization of Atg9 and Lamp1, which is a lysosomal protein that would normally not be near Atg9 vesicles. These cells were incubated with pepstatin and E64d to inhibit lysosomal degradation of Atg9. The results show dramatically increased colocalization of Atg9 with LAMP1 in Dnm2 -/- cells after induction with CCCP (Fig 7I and J).

Lastly, we used Western blots to monitor Atg9 levels in Dnm2 -/- cells. Levels of Atg9 protein decrease when mitophagy is induced in Dnm2 -/- cells with CCCP but not in wild type cells (Fig 7K). This decrease is halted with Bafilomycin, suggesting that Atg9 is consumed by lysosomal degradation in Dnm2 -/- cells (Fig 7I and J), but not in wildtype cells. Together, these experiments show that Atg9 retrieval from phagophores is impaired in Dnm2 -/- cells and that Atg9 is subsequently degraded in lysosomes. Although the overall rates of autophagy do not decrease in Dnm2 -/- cells there may be other functional consequences because some of the early autophagy proteins such as Atg9 and SNAREs will be turned over more rapidly than in wildtype cells and their retention on phagophores could interfere with downstream processes.

In sum, our data suggest a model with two stages that are affected by Dnm2 and EndoB1. In the first stage, EndoB1 and Dnm2 act together to promote retrieval of Atg9 (Fig 7L). At this stage, Dnm2 prevents further progression along the autophagy pathway until the phagophore is fully grown. In the second stage, EndoB1 relieves the inhibitory function of Dnm2, thus allowing further progression along the pathway with fusion to Rab7-containing late endosomes and lysosomes and completion of the autophagy process (Fig 7M).

## Discussion

Our data support two unexpected roles for Dnm2 during autophagy. Dnm2 promotes the retrieval of Atg9 and other membrane proteins from growing phagophores and it acts as a negative regulator of downstream fusion events, thus providing a mechanism for coupling the orderly completion of phagophores to a switch from early to late stages of autophagy (Fig 7L and M). Retrieval of Atg9 from phagophores could be inferred from the previous finding that Atg9 is present on incoming vesicles, but absent from mature autophagosomes (Orsi et al., 2012). We now provide five lines of evidence showing that Dnm2 is required for this retrieval process and for the retrieval of other transmembrane proteins from phagophores.

First, we detected substantial colocalization of Atg9 with LC3 in Dnm2 -/- cells and in extracts when autophagy is induced. These results suggest that transmembrane proteins in Atg9 vesicles colocalize with LC3 containing membranes when Dnm2 function is impaired. Similar effects were observed with other methods to impair Dnm2 function, such as Dnm2 siRNA and chemical inhibitors of Dnm2 function. Second, the numbers of Atg9 vesicles decreased when autophagy was induced in Dnm2 -/- cells, suggesting that Atg9 vesicles fail to reform after fusion with phagophores. Third, the levels of Atg9 protein were reduced in Dnm2 -/- cells and they were further reduced when autophagy was induced. This reduction is blocked by Bafilomycin, suggesting that Atg9 is incorporated in autophagosomes and degraded by autophagy when Dnm2 function is impaired. Fourth, SNARE proteins needed for fusion of Atg9 vesicles with phagophores mix with LC3 when autophagy is induced in Dnm2 -/- cells. Fifth, Dnm2 colocalizes with Atg9, EndoB1 and LC3 when autophagy is induced, as shown with a variety of different methods. Taken together, these data show that Dnm2 is required for recycling transmembrane proteins from phagophores.

It was previously shown that Dnm2 promotes the budding of Atg9 vesicles from recycling endosomes, either in concert with EndoB1 (Takahashi et al., 2016) or with SNX-18 (Soreng et al., 2018). It was concluded that the Atg9 vesicles, which are formed at recycling endosomes, travel to phagophores at other sites in the cell where they deposit their membrane. In this scenario, Dnm2 is needed for the delivery of membrane to phagophores, but our data shows that autophagy proceeds unimpeded in Dnm2 -/- cells. A possible solution for these seemingly conflicting results comes from a recent study in which it was shown that phagophores start off as recycling endosomes (Puri et al., 2018). Our results agree with the possibility that Dnm2 helps form Atg9 vesicles from recycling endosomes, but only when those endosomes are being transformed into phagophores. The Dnm2-mediated formation of Atg9 vesicles is therefore not needed for phagophore growth. Instead, these vesicles retrieve Atg9 (and possibly other proteins) from growing phagophores.

The triggers for Dnm2 relocation to growing phagophores have not yet been determined. Dnm2 normally binds to PI(4,5)P_2_ on the plasma membrane, but PI(4,5)P_2_ is also found on internal membrane compartments, such as endosomes (Tan, Thapa et al., 2015). Our experiments with the PI(4,5)P_2_ reporter PLC-delta PH domain suggest that Dnm2 -/- cells accumulate this lipid on autophagic membranes, consistent with a role in the recruitment of Dnm2 to phagophores. Interestingly, Atg9 was recently shown to bind yet another BAR domain protein, Arfaptin 2, which in turn delivers a PI(4)P-kinase to autophagosomes (Judith, Jefferies et al., 2019). This PI(4)P-kinase could direct the formation of PI(4,5)P_2_ if augmented by a PI(4)P-5-kinase. Dynamins can, however, also bind to PI(3,5)P_2_, as shown with biochemical studies (Yarar et al., 2008). Phagophores have ample supplies of PI(3)P and there are reports of PI(3,5)P_2_ generated by PIKfyve from PI(3)P on autophagic membranes (Martin, Harper et al., 2013) suggesting PI(3,5)P_2_ as an alternative signal for Dnm2 localization to autophagic membranes.

Other aspects of the fission process have also not yet been characterized, but some parallels can be drawn with Dnm2-mediated fission events at the plasma membrane. Along with the well-known roles of Dnm2 in clathrin- and caveolin-dependent endocytosis (Henley, Krueger et al., 1998, van der Bliek, Redelmeier et al., 1993) as well as in clathrin/caveolin-independent endocytosis (Conner & Schmid, 2003), Dnm2 also contributes to fast endocytosis (FEME), where this protein acts in concert with EndoA1 (Renard et al., 2015), and it contributes to kiss-and-run fission and fusion, which does not require additional vesicle budding proteins (Alabi & Tsien, 2013, Fulop, Doreian et al., 2008). These last two processes can be relevant paradigms for the actions of Dnm2 during autophagy; EndoA1 is very similar to EndoB1, which we show participates with Dnm2 in autophagy, while kiss-and-run fission and fusion can mediate lipid transfer through a transient fusion pore without also transferring proteins (Monck, Alvarez de Toledo et al., 1990). The effects of the dynamin activator Ryngo 1-23 and inhibitor Dynole 34-2 on the appearance of Dnm2 spots sandwiched between Atg9 and LC3 containing membranes (Fig 6B) are similar to the effects of these chemicals on dynamin-mediated kiss-and-run events at the plasma membrane (Jackson et al., 2015, Lasic et al., 2017). Together, these results suggest a speculative model in which scission at autophagosomes is mediated by the concerted actions of Dnm2 and EndoB1, allowing for retrieval of Atg9 (Fig EV5).

The role of Dnm2 as an inhibitor of autophagy, preventing the transition from phagophore to autophagosome when EndoB1 is mutated, was uncovered by the suppression of the EndoB1 mutant phenotype when Dnm2 was also mutated in DKO cells. This novel role provides a switch between Atg9-mediated growth of the phagophore and later stages in which growth stops and the phagophore fuses to late endosomes and lysosomes. This switch is evident with Rapamycin and CCCP-induced autophagy, because EndoB1 -/- cells accumulate LC3-II, while the DKO cells do not, but it was not observed with starvation-induced autophagy, possibly resulting from alternative requirements for EndoB1 or alternative sites for phagophores (near ERGIC or MAM) (Ge, Melville et al., 2013, Ge, Zhang et al., 2017, Graef, Friedman et al., 2013, Hamasaki, Furuta et al., 2013). Inhibition does, nevertheless, provide an elegant mechanism for coupling the transition from early to late stages of autophagy induced by CCCP or Rapamycin.

In conclusion, our results have helped define two novel functions for Dnm2 during autophagy. The proposed roles of Dnm2 and EndoB1 as the membrane scission proteins that promote retrieval of Atg9 from phagophores is supported by colocalization and functional studies with gene deletions, siRNA and chemical inhibitors. The newly uncovered regulatory pathway controlling the transition between early and late stage of autophagy suggests additional interactions between EndoB1, Dnm2 and downstream factors affecting fusion to late endosomes and lysosomes. Future studies should help clarify this additional role of Dnm2 and EndoB1.

## Materials and methods

### Plasmids

The pMito-DsRed2 plasmid was from Clontech. Addgene provided pcDNA3-mRuby2 (#40260), GFP-C1-PLCdelta-PH (#21179), Dnm2-EGFP (#34686), Dnm2-mCherryN1 (#27689), ptfLC3 (#21074), pMXs-puro-RFP-ATG9A (#60609), pEGFP-LC3 (# 24920), GFP-Rab7A (#61803), mTaqBFP2-ER-5 (#55294), pcDNA3-mRuby2 (#40260), GFP-Atg2 (#36456), GFP-Fip200 (#38192), GFP-VAMP7 (#42316) and pmRFP-LC3 (#21073). Yu Sun and Lars Dreier (Dept. of Biological Chemistry, UCLA) provided the GFP-Stx17 plasmid. F. X. Soriano (Department of Cell Biology, University of Barcelona) provided the GFP-Bax plasmid. E. C. Dell’Angelica (Department of Human Genetics, UCLA School of Medicine) provided the GFP-Lamp1 plasmid. To generate EndophilinB1-mRuby2, EndophilinB1 coding sequences from cDNA were fused in frame to mRuby2 by PCR cloning into pcDNA3-mRuby2. EGFP-Atg9, Dnm2-RFP and mTaqBFP2-LC3 constructs were generated in the same way with coding sequences from pMXs-puro-RFP-ATG9A, Dnm2-EGFP and mTaqBFP2-ER-5, respectively.

### Cell culture, transfection, gene knockout and chemical treatments

WT HeLa cells were from James Wohlschlegel (Dept. of Biological Chemistry, UCLA) and WT MEFs were from David Chan (Dept of Biology, CalTech). All cell lines were periodically checked for Mycoplasm. MEFs were grown in DMEM with 10% FBS. Transient transfections were done with jetPRIME following manufacturer’s instructions (Polyplus). For siRNA, cells were grown in 6cm dishes, transfected with 50nM oligonucleotides using RNAimax (Invitrogen) and analyzed 72h later. MEF cells with conditional knockout of all three dynamin isoforms (TKO) were obtained from P. De Camilli (Yale School of Medicine, New Haven). Growth and conditional knockout procedures for these cells were as described (Ferguson, Raimondi et al., 2009, Park et al., 2013, Raimondi, Ferguson et al., 2011). Deletions in Dnm2, EndoB1 and Drp1 genes were introduced in MEF cells using CRISPR/Cas9 methods, as described (Ran, Hsu et al., 2013). Target sites were identified using the Boutros Lab Website (http://www.e-crisp.org/E-CRISP/designcrispr.html). Dnm2 gRNA was: 5’-GATGGCAAACACGTGCTTGT-3’. EndoB1 gRNAs were: 5’-GCAGGAACTGAGTTTGGCCC-3’; and 5’-GGATTTCAACGTGAAGAAGC-3’. Drp1 gRNA was: 5’-GCAGTGGGAAGAGCTCAGTGC-3’. The gRNAs were cloned in the px459 plasmid, containing puromycin resistance and Cas9 genes, and 1.2μg was transfected into MEF cells, followed by 24hr puromycin selection. Surviving colonies were isolated, genotyped and analyzed with western blots. HeLa cells were also grown in DMEM with 10% FBS. The gRNA used to knock out Dnm2 in HeLa cells was: 5’-ACAGGGGCCGGCCCTGGACC-3’. The knock out procedure was as described for MEF cells.

Amino acid starvation was induced by washing cells twice in EBSS with calcium and magnesium (Thermo Fisher Scientific), and incubated for the indicated length of time. The following chemicals with their final concentrations were from Sigma-Aldrich: 20μM CCCP, 40μg/ml antimycin A, 10μM ionomycin, 10μM actinomycin D and 1μM 4-Hydroxy Tamoxifen. The following chemicals with their final concentrations were from InvivoGen: 0.5μM Bafilomycin A1, 10μM rapamycin and 1μM YM-201636. Staurosporine (Thermo Fisher Scientific) was used at 2μM. Z-VAD-FMK (BD Bioscience) was used at 20μM. Dynasore (EMD Millipore) was used at 80μM. VPS34-IN1 (Selleckchem) was used at 10μM.

### Immunoblotting, immunofluorescence, and proximity ligation assay

Total cell lysates for Western blots were made with RIPA buffer. Samples were subjected to SDS-PAGE, transferred to nitrocellulose or PVDF membranes, blocked with Odyssey Blocking Buffer (LI-COR), and incubated overnight at 4°C with primary antibodies. Membranes were then washed with PBS-T and incubated with IRDye 800CW or 670RD secondary antibodies (LI-COR). Fluorescent bands were detected with an Odyssey scanner and analyzed with Image Studio Software (LI-COR). For immunofluorescence, cells were grown on 12mm coverslips, fixed for 10min with 4% paraformaldehyde in PBS, and permeabilized for 5min with 0.1% Triton X-100 in PBS, blocked for 1h with Goat or Donkey Serum in PBS-T and incubated with primary antibodies. For immunofluorescence of Dnm2, cells were first permeabilized with 0.05% digitonin to wash out soluble cytosolic proteins (Liu et al., 1998). Secondary antibodies were Alexa Fluor 488-, 594- or 647-conjugated goat anti-mouse or rabbit IgG (Invitrogen). Proximity ligation assays were conducted with Duolink as recommended by the manufacturer (Sigma-Aldrich).

### Antibodies

Rabbit anti-Endophilin A1, mouse anti-SQSTM1 (p62), rabbit anti-dynamin 1, rabbit anti-dynamin 2, mouse anti-Hsp60, rabbit anti-cytochrome c and mouse anti-VDAC1/Porin were from Abcam. Rabbit anti-LC3B and mouse anti-tubulin antibodies were from Sigma Aldrich. Goat anti-dynamin 2 and rabbit anti-Tom20 were from Santa Cruz Biotechnology. Mouse anti-Tim23 and mouse anti-Drp1 were from BD Biosciences. Rabbit anti-OPTN and rabbit anti-RAB7A were from Proteintech. Mouse anti-Endophilin B1 was from Imgenex. Rabbit anti-SNX18 and rabbit anti-ATG9A were from GeneTex.

### Fluorescence microscopy

For live imaging, cells were grown in glass bottom dishes (MatTek). Cells were viewed with a Marianas spinning disc confocal from Intelligent Imaging, which uses an Axiovert microscope (Carl Zeiss Microscopy) with a CSU22 spinning disk (Yokogawa), an Evolve 512 EMCCD camera (Photometrics) and a temperature unit (Okolab). Cells were imaged at 37°C with 40x/1.4 and 100x/1.4 oil objectives. Super resolution images were acquired with SIM using a DeltaVision OMX SR (General Electric). Fiji ImageJ software was used for image analysis. Time lapse images were acquired with a Zeiss Airyscan microscope equipped with a Fast-Airyscan module and processed with Airyscan software.

### Subcellular fractionation

Extracts for imaging of partially purified autophagosomes were essentially made as described (Gao et al., 2010b). In brief, MEF cells were transfected with EGFP-Atg9 and RFP-LC3. After 16h these cells were treated for by 4hr with 20 μM CCCP and 0.5 μM Bafilomycin. Cells were harvested in mitochondrial isolation buffer (0.25 M sucrose, 1 mM EDTA, 20mM HEPES, pH7.4) on ice, disrupted by passing 20 times through a 22 gauge needle. Nuclei and unbroken cells were removed with low speed centrifugation (10 min at 600g) followed by enrichment of autophagosomes with medium speed centrifugation (20 min at 10,000g) and washing twice in 1.0 M NaCl in mitochondrial isolation buffer with protease inhibitor to dissociate clusters of vesicles that were potentially held together by ionic interactions. Samples of the resulting fractions were mounted on slides and examined by fluorescence microscopy.

For Ficoll gradients, WT and Dnm2 -/- MEF cells were treated for 4 hr with 0.5mM Bafilomycin A, followed by 2 hr with 0.5mM Bafilomycin A and 5 μM Rapamycin to induce autophagy. Cells were then fractionated as described (Orsi et al., 2012), but with a few modifications. In brief, cells from 4 confluent 15 cm dishes were washed, scraped and resuspended in homogenization buffer (250 mM sucrose, 20 mM HEPES/KOH pH 7.4, 0.5 mM PMSF), followed by homogenization with 15 passes through a 25 G needle. Nuclei and unbroken cells were removed with low speed centrifugation. The equivalent of 2 mg protein was loaded on a 1-22% Ficoll gradient with a total volume of 10 ml layered on a 1 ml 45% Nycodenz cushion in homogenization buffer. These preparations were centrifuged in a Beckmann SW41 rotor for 14.5 hr at 20 krpm at 4°C. Twenty fractions of 0.5 ml were collected and even numbered fractions were analyzed with Western blots of 40 μl samples.

## Acknowledgements

We are grateful for many helpful discussions and sharing of reagents with Lars Dreier, Yu (Sammy) Sun at UCLA. CMK was supported by NIH grants R01GM61721, R01GM073981 and R01DK101780. AMvdB was supported by NIH grant U01GM109764.

## Author Contributions

AMR conducted experiments and helped design them. CIM, SI, JS and BR conducted experiments. CMK was a co-supervisor and helped design the experiments. AMvdB supervised the project, helped design the experiments and wrote the paper.

## Conflict of interest

The authors declare that they have no conflict of interest.

## Figure legends

**Figure EV1.**
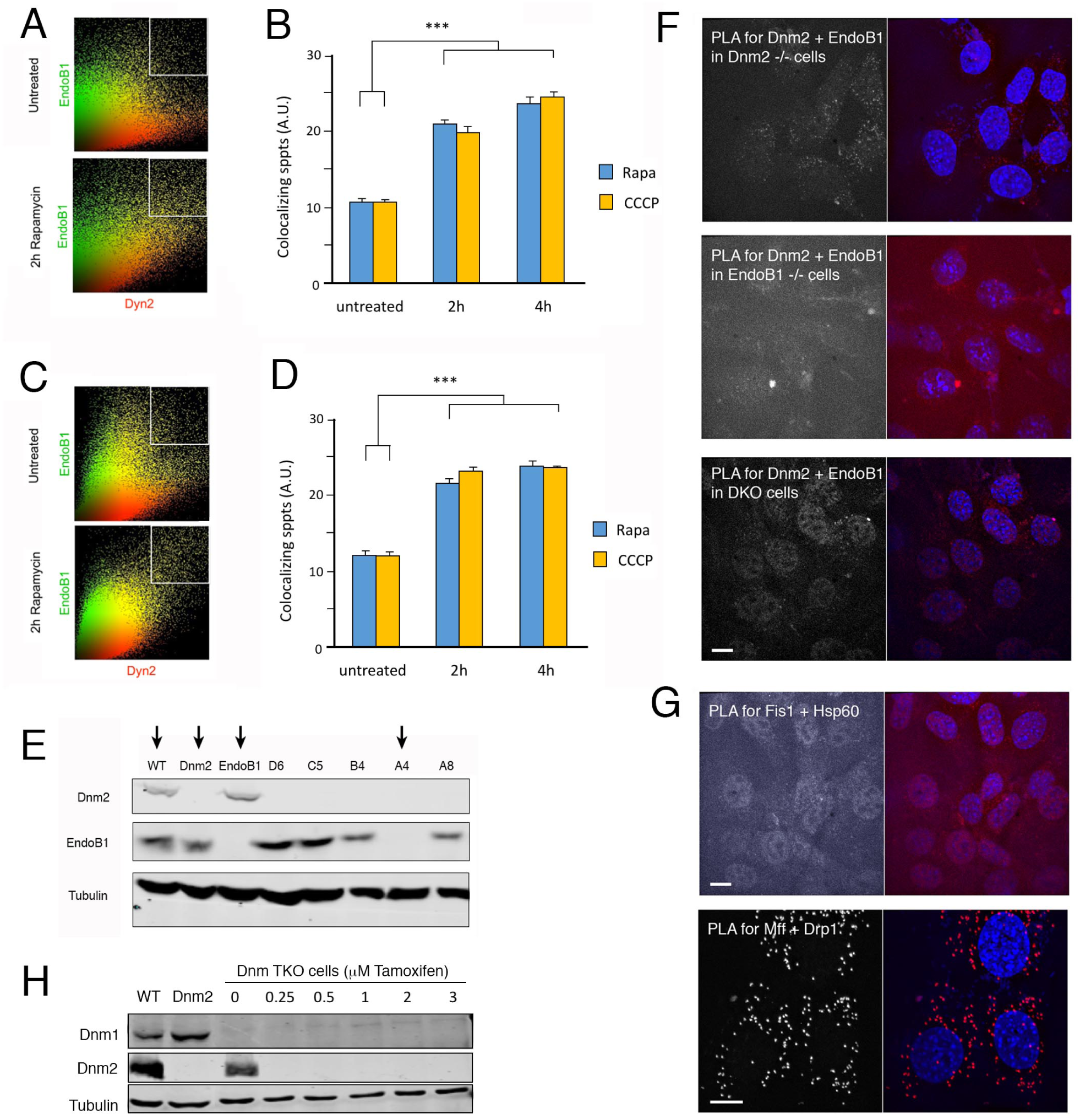
Colocalization of Dnm2 and EndoB1 shown with scatterplots. A Scatterplots of immunofluorescence images of cells treated for 2h with or without rapamycin to induce autophagy. B Fraction of spots in the colocalizing quadrants of the scatterplots, shown before and after 2 or 4 hr treatment with rapamycin or CCCP. (SE and unpaired Student’s t-test, n= 3). C, D As in panels A and B, but with transiently expressed GFP- and RFP-tagged proteins. E Western blots showing derivation of Dnm2, EndoB1 and DKO cells. Dnm2 and EndoB1 -/- cells were generated by CRISPR/Cas9 in MEFs. DKO cells were generated by additional knockout of EndoB1 in Dnm2 -/- cells. Clone A4 was chosen for further studies. F PLA signals for Dnm2 and EndoB1 interactions were absent from cells with mutations in one or both of these proteins. G As negative control for PLA, cells were incubated with Fis1 and Hsp60 antibodies, which detect proteins that are on the outside or inside of mitochondria and are therefore too far apart for generating PLA signals. As positive control, cells were incubated with antibodies for Drp1 and Mff, which detect well-characterized binding partners. Scale bar is 10 μm. H Western blots of conditional dynamin knockout cells. Cre-recombinase was induced with tamoxifen at the indicated concentrations for 24 hr and cells were grown for an additional 4 days with fresh medium before western blot analysis of Dnm1 and Dnm2 expression. Although Dnm1 is not normally expressed in these cells and Dnm3 is undetectable with currently available antibodies (Park et al., 2013), all three genes were floxed with 3 μM tamoxifen to ensure complete lack of dynamins.

**Figure EV2.**
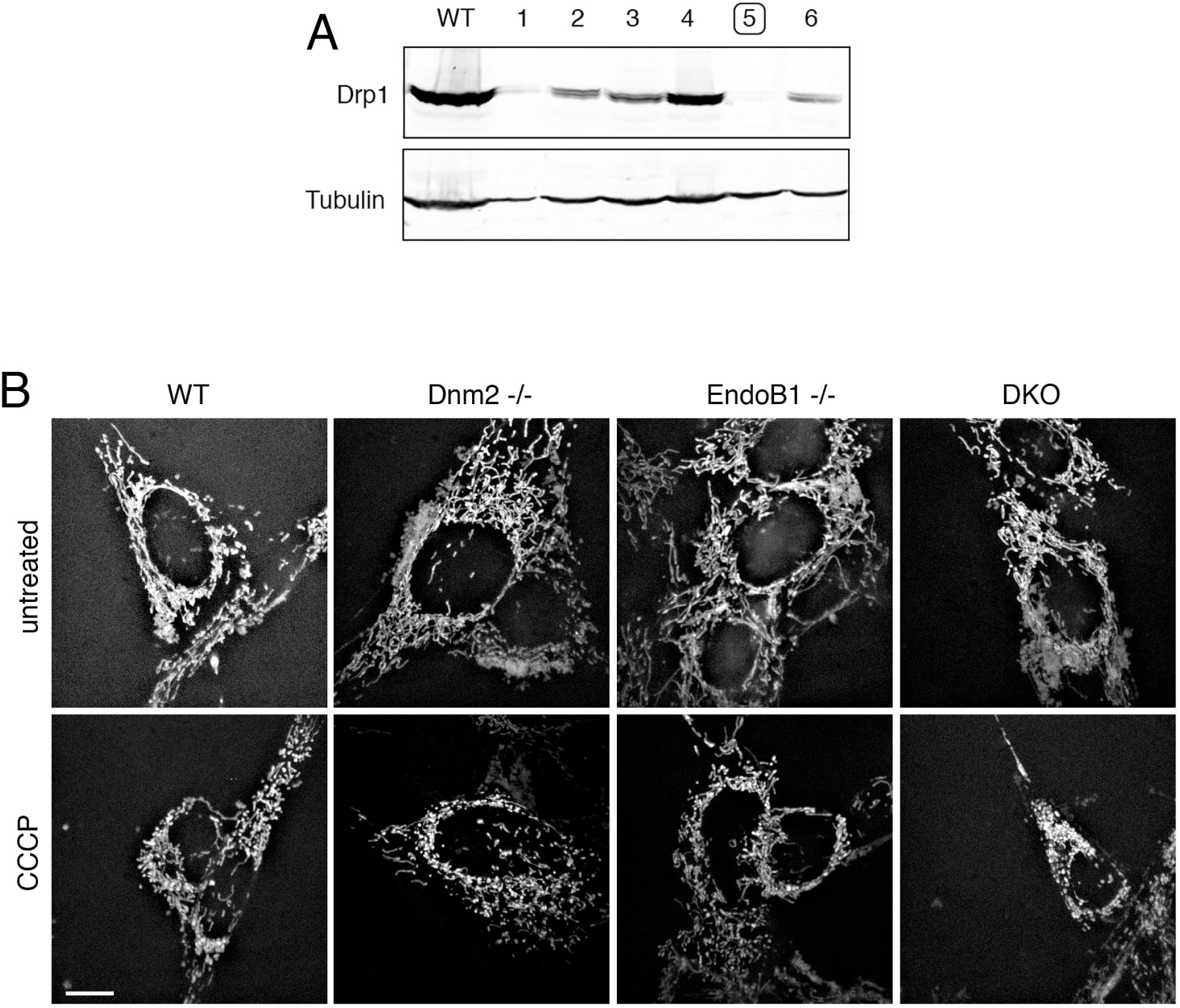
Western blots of Drp1 deletion mutants and effects of Dnm2 and EndoB1 mutations on mitochondrial fission shown with live cells. A Drp1 deletions mutants were generated with CRISPR/Cas9 technology. Clone number 5 was selected for further analysis. B The images show MEFs transfected with mitochondrial DsRed and genotypes as indicated. Cells were treated with or without 20 μM CCCP for 30min. Scale bar is 10 μm.

**Figure EV3.**
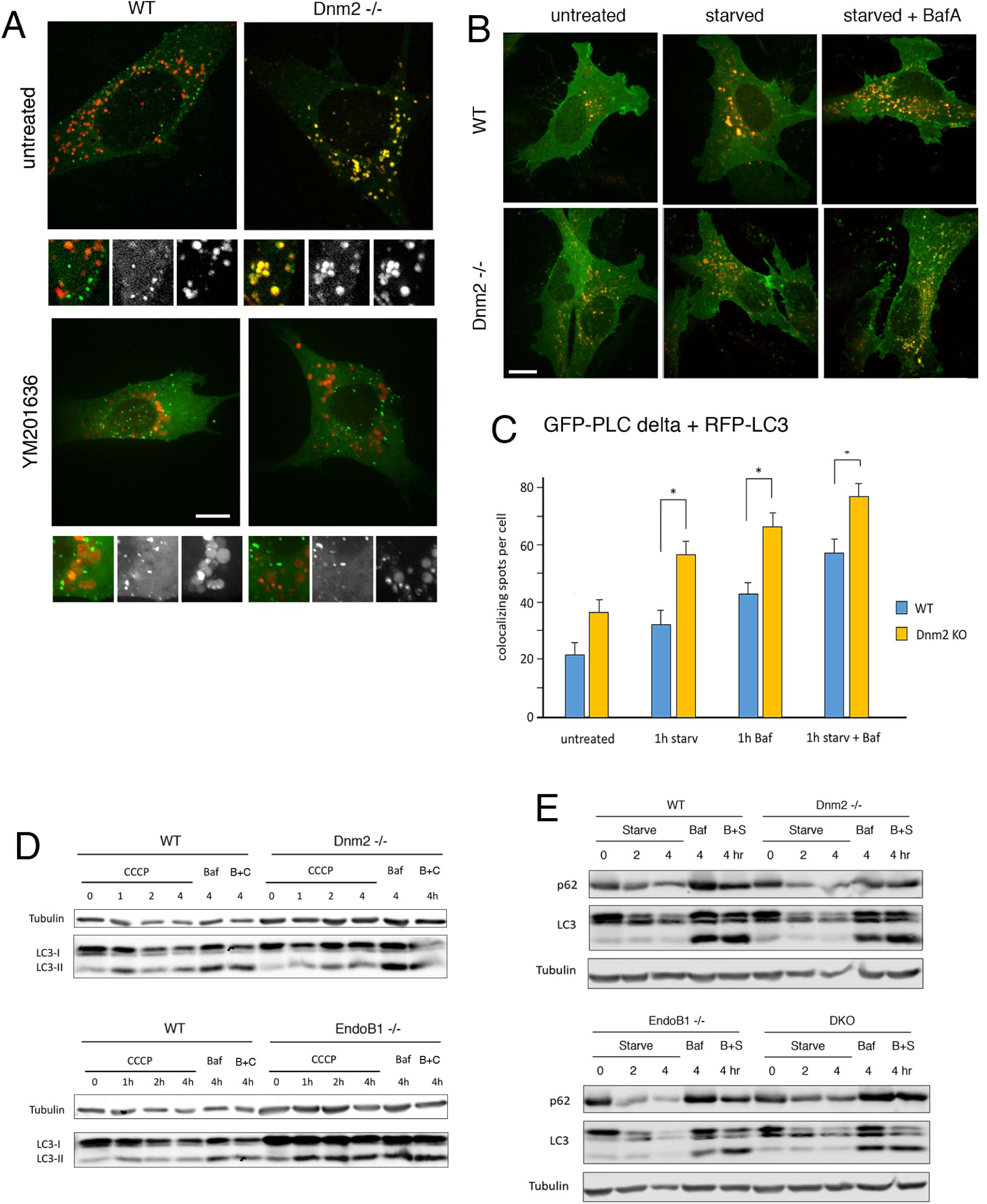
Altered distributions of PIP_2_ in Dnm2 -/- cells and levels of autophagy proteins in Dnm2 and EndoB1 -/- cells. A Overexpressed GFP-Dnm2 spontaneously colocalizes with RFP-LC3 in Dnm2 -/- cells, but not in WT cells. This colocalization is disrupted by the PIKfyve inhibitor YM201636. Enlarged portions of the cells (2X) are shown below each image with merged, Dnm2 (green) and LC3 (red) channels. B Colocalization of GFP fused to the PH domain of PLC delta with RFP-LC3 is also observed in Dnm2 -/- cells. C Histograms show increased colocalization of the GFP-tagged PH domain of PLC delta with RFP-LC3 in Dnm2 -/- cells. For the histograms, 50 cells per experiment, SEM, n=3, unpaired Student’s t-test. Scale bar is 10 μm. D Example of Dnm2 and EndoB1 -/- effects on CCCP induced LC3 lipidation. There is an accumulation of LC3-II in EndoB1 -/- cells but not in Dnm2 -/- cells. E Starvation induced autophagy is not affected by mutations in Dnm2 or EndoB1 as shown with p62 turnover and LC3 lipidation.

**Figure EV4.**
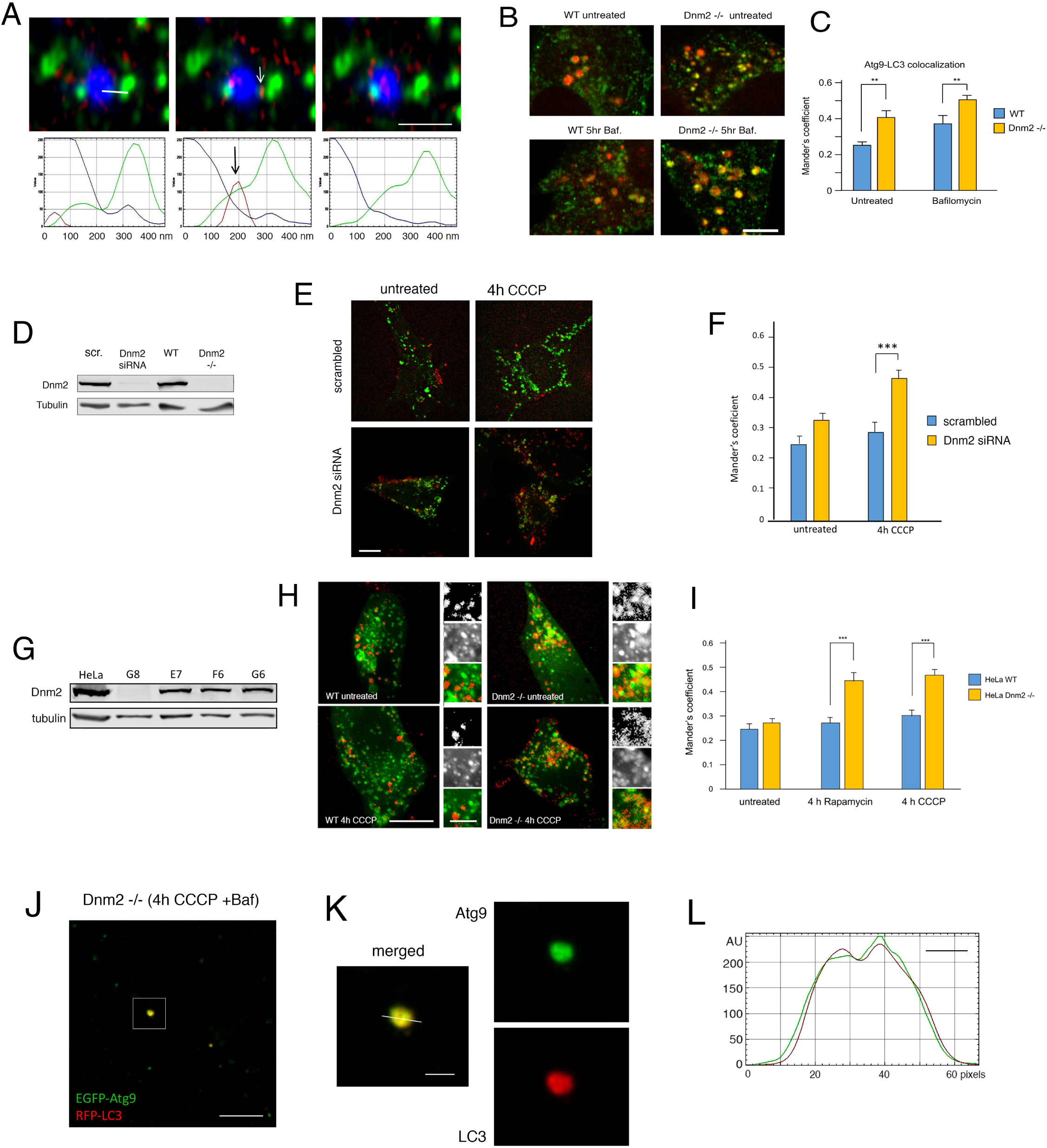
Atg9 localization in live MEF cells, in Dnm2 siRNA MEF cells and in Dnm2 -/- HeLa cells. A Example of a Dnm2 spot transiently wedged between Atg9 and LC3 containing organelles. Time lapse images of GFP-Atg9, RFP-Dnm2 and BFP-LC3 were taken at 350 msec intervals. An arrow in the middle panel points to a potential scission event. The bar in the first panel shows the line used for the intensity tracings shown in the bottom panels. Scale bar is 1 μm. B, C Effects of Dnm2 on Atg9 retrieval in live cells. Images of BFP-LC3 (shown in red) and GFP-Atg9 in live WT and Dnm2 -/- cells, untreated or after 5 hr with Bafilomycin. Scale bar is 5 μm. Histograms showing increased colocalization of BFP-LC3 and GFP-Atg9 in Dnm2 -/- cells with or without Bafilomycin treatments. 50 cells per condition in each experiment, n=3, SE and unpaired student’s T test. D Western blot showing depletion of Dnm2 after transfection of WT cells with siRNA oligonucleotides. E RFP-Atg9 colocalizes with EGFP-LC3 after 4h treatment with CCCP in Dnm2 siRNA cells, but not in mock-transfected cells. F Histogram showing increased colocalization of RFP-ATG9 and GFP-LC3 in CCCP-treated Dnm2 siRNA cells. Scale bar is 10 μm. G Western blot showing depletion of Dnm2 in a CRISPR/Cas9 generated clone HeLa cells. Clone G8 was chosen for further analysis. H RFP-Atg9 colocalizes with EGFP-LC3 in Dnm2 -/- HeLa cells treated with CCCP, but not in wild type cells. Scale bar is 10 μm for whole cells and 3 μm for enlarged portions. I Histogram showing increased colocalization of RFP-ATG9 and GFP-LC3 in Dnm2 -/- cells after incubation with rapamycin or CCCP. J Image of autophagosome containing fraction from Dnm2 -/- cells transfected with EGFP-Atg9 and RFP-LC3 followed by 4hr treatments with CCCP and Bafilomycin to induce mitophagy and to accumulate autophagosomes. Autophagosomes were enriched by differential centrifugation. Scale bar is 10 μm. K Enlargements show an individual spot with the line used to plot intensities. Scale bar is 2 μm. The x-axis in the fluorescence intensity plot shows numbers of pixels (24 pixel/μm) and the bar shows 500 nm. L Line tracings show fluorescence intensities

**Figure EV5.**
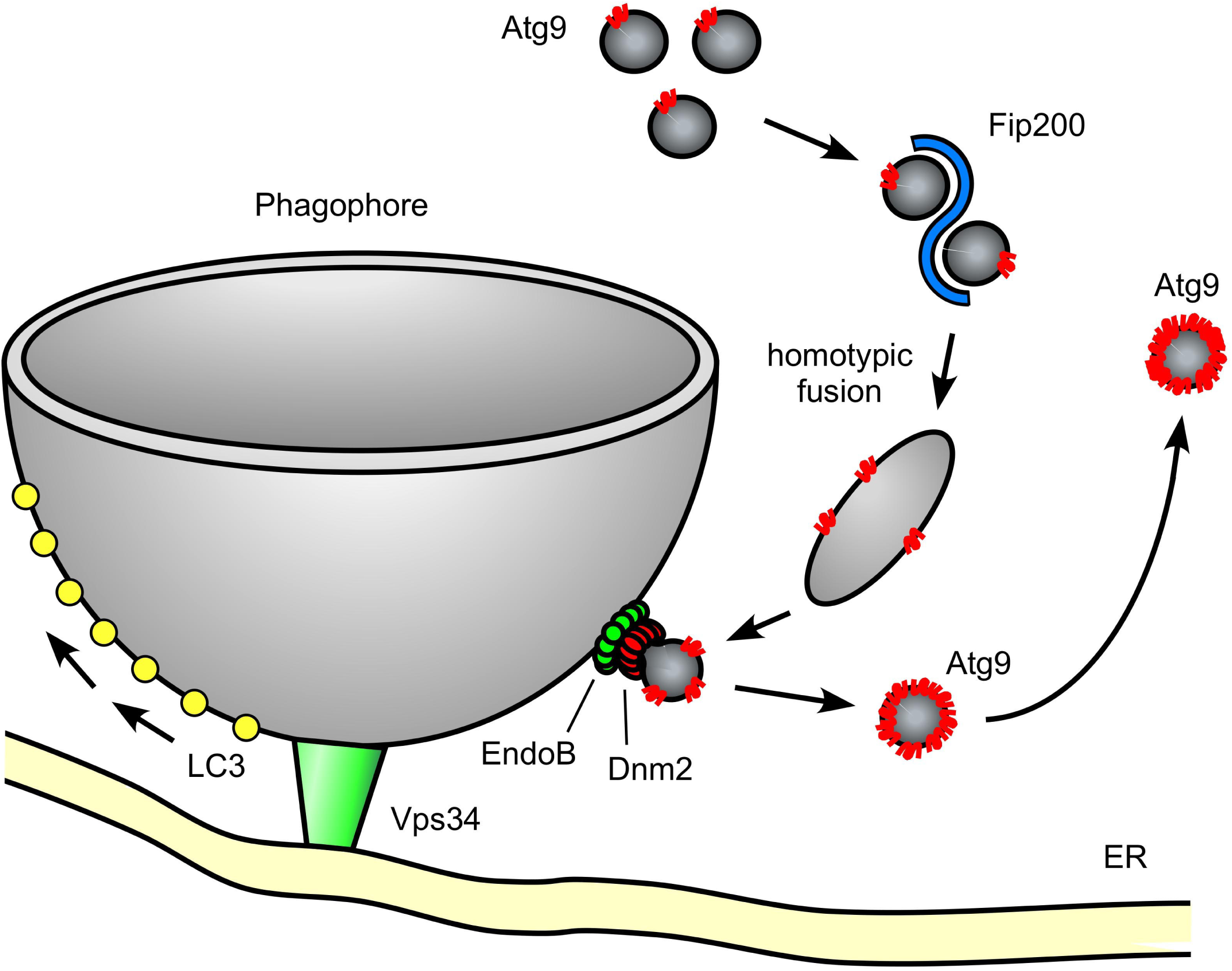
Model Dnm2 and EndoB1 functions during the addition of membrane to a growing phagophore. Atg9 vesicles derived from other compartments, such as the Golgi and early endosomes collect near the site of autophagy, where they undergo homotypic fusion assisted by Fip200. These larger Atg9 containing structures can then fuse to the growing phagophore. After depositing membrane, Atg9 is retrieved by Dnm2 and EndoB1 mediated scission and returned to the originating organelles.

